# Tumor-Intrinsic IFNγ Signaling and Niche Adaptation Drive Early Colonization in Ovarian Cancer Metastasis

**DOI:** 10.1101/2025.08.13.669778

**Authors:** Emilija Aleksandrovic, Shaneann R. Fross, Samantha M. Golomb, Wei Ma, Xiyu Liu, Zhuo Zhao, Nikitha M. Das, Tanner C. Reese, Hazel M. Borges, Ates Tenekeci, Lei Yu, Jacqueline Lopez, Min Zhao, Zhenyu Zhong, Kevin M. Dean, Jayanthi Lea, Lin Xu, M. Sharon Stack, Siyuan Zhang

**Affiliations:** Department of Pathology, UT Southwestern Medical Center, Dallas, TX, 75390, USA; Harold C. Simmons Comprehensive Cancer Center, Dallas, TX, 75235, USA; Department of Urology, UT Southwestern Medical Center, Dallas, TX, 75390, USA; Lyda Hill Department of Bioinformatics, UT Southwestern Medical Center, Dallas, TX, 75390, USA; Cecil H. And Ida Green Center for Systems Biology, UT Southwestern Medical Center, Dallas, TX, 75390, USA; Quantitative Biomedical Research Center, Department of Health Data Science & Biostatistics, Peter O’Donnell Jr. School of Public Health, UT Southwestern Medical Center, Dallas, TX, 75390, USA; Department of Pediatrics, UT Southwestern Medical Center, Dallas, TX, 75390, USA; Department of Biological Sciences, College of Science, University of Notre Dame, Notre Dame, IN 46556, USA; Department of Immunology, UT Southwestern Medical Center, Dallas, TX, 75390, USA; Department of Obstetrics and Gynecology, UT Southwestern Medical Center, Dallas, TX, 75390, USA; Department of Chemistry and Biochemistry, University of Notre Dame, Notre Dame, IN 46556, USA; Mike and Josie Harper Cancer Research Institute, University of Notre Dame, 1234 N. Notre Dame Avenue, South Bend, IN 46617, USA

**Author notes:** **Correspondence to:** Siyuan Zhang, M.D., Ph.D., Department of Pathology, UT Southwestern Medical Center, ND6.200A, 6001 Forest Park Rd., Dallas, TX 75235.

**Keywords:** Ovarian cancer, Metastasis, Disseminating Tumor Cells (DTCs), Clonal dynamics, Metastatic niches, IFN-gamma (IFNγ), Tumor-associated macrophages (TAMs), Anoikis, Single-cell RNA sequencing, MetTag barcoding, Clonal tracing, In vivo CRISPR Screen

## Abstract

Metastasis is an emergent continuum driven by evolving reciprocal adaptations between disseminating tumor cells (DTCs) and specialized niches of different organs. The interplay between intrinsic and niche-driven mechanisms that enable DTCs to survive and home to distant organs in peritoneal ovarian cancer metastasis remains incompletely understood. Here, we present MetTag, a single-cell barcoding and transcriptome profiling approach with single cell clonality barcodes and time-stamped batch identifiers (BC.IDs) to resolve metastasis clonality and temporal dynamics of DTC colonization. Deep sequencing of MetTag barcodes revealed enrichment of early-disseminated clones across metastatic sites, and targeted depletion of pioneer DTCs diminished the outgrowth of subsequent arriving DTCs. Subsequent MetTag-coupled single cell RNA sequencing (scRNA-seq) on ascites and metastasis-bearing omenta revealed a distinct interferon-gamma (IFNγ)-centric transcriptional trajectory selectively enriched among pioneer clones. *In vivo* CRISPR/Cas9 screening of niche-specific signatures demonstrated that the tumor-intrinsic IFNγ response is functionally required for peritoneal metastasis. Knockout of the IFNγ receptor 1 (*Ifngr1*) in the initial pioneer DTCs significantly reduced total metastatic burden, revealing a critical window of time in which IFNγ signaling shapes the post-seeding metastatic niche (PSMN) and subsequent metastatic evolution. Mechanistically, the tumor-intrinsic IFNγ response induced Poly(ADP-Ribose) polymerase family member 14 (Parp14) and peritoneal macrophages cooperatively shield DTCs from anoikis by promoting pro-survival signaling. Our study defines the temporal clonal architecture of peritoneal niches and reveals a “first come, first served” adaptation principle, where pioneer colonizer fitness determines the success of subsequent colonizers.

## INTRODUCTION

Cancer metastasis encompasses continuous metastatic dissemination from the primary tumor, accompanied by passive clonal selection, and active epigenetic adaptation in response to the metastatic niche^1^. Resolving the clonal, temporal, and microenvironmental determinants of this process remains a significant challenge in the field. Unlike many cancer types where hematogenous metastasis predominates, high-grade serous ovarian carcinoma (HGSOC) exhibits a unique pattern of transcoelomic, local spread within the peritoneal cavity^2^. In HGSOC, malignant ascites functions as a dynamic “reservoir” of free-floating cancer cells and cell clusters – DTCs detached from the primary tumor – initially seeding locally and colonizing the greater omentum and other peritoneal organs^3^. Different from immunologically “cold” tumor microenvironments, malignant ascites is heavily enriched with immune cells and acellular factors known to influence tumor progression. Consequently, ovarian cancer DTCs - continuously shed from the primary tumor - must overcome immune surveillance and adapt to the unique intraperitoneal microenvironment to establish colonization ^4,5,6,7,8^. In contrast to the pre-metastatic niche (PMN), which describes systemic conditioning of distant sites by the primary tumor prior to DTC arrival^9,10^, pioneer and subsequent DTC colonization actively remodel the metastatic niche, forming a progressively co-evolving post-seeding metastatic niche (PSMN) that influences the ultimate outcome of metastasis. Clinical studies in multiple tumor types, including HGSOC, have corroborated the presence of DTCs in bone marrow and distant organs of patients well before reaching advanced (stage III & IV) tumor stages^11–17^. Despite early metastatic dissemination observations, underlying molecular mechanisms that modulate reciprocal interplay between DTC seeding waves and PSMN evolution driving metastasis outgrowth remain largely unknown.

Here, we conducted comprehensive clonal tracing, single-cell RNA sequencing (scRNA-seq), *in vivo* CRISPR screening, and Cre-based niche-tracing analysis to dissect the complex interplay between continuously arriving DTCs and ovarian cancer PSMN. We identified a critical pioneer DTC dependence on a niche-adaptive IFNγ response signature which favors early pioneer DTC outgrowth. These IFNγ-responsive pioneer DTCs dictate the success of later arriving DTCs. We further demonstrate that DTC-PSMN co-conditioning enriches for a macrophage population capable of protecting DTCs from anoikis-mediated death via paracrine signaling. Together, our data reveal a “first come, first served” principle where pioneer DTC clones dictate the fate of later DTCs within a relatively short and defined temporal window.

## RESULTS

### MetTag Captures Spatiotemporal Clonal Dissemination Patterns in Ovarian Metastasis

To capture the transcriptional and clonal dynamics, we employed an RNA-level barcoding system based on the LARRY (**<u>L</u>**ineage **<u>A</u>**nd **<u>R</u>**NA **<u>R</u>**ecover**<u>y</u>**) barcoding construct, which enables single-cell transcriptome-level lineage tracing^18^. Critically, we added a unique batch identifier “BC.ID” as a time stamp for distinguishing time points (TPs) representing early or later-arriving DTCs. Within the U6-driven barcode sequence, a Chromium 10x capture sequence 1 (CS1) allows for simultaneous capture of the LARRY barcode transcript with mRNA transcripts (**Fig.1A & S1A**). LARRY barcode diversity in each resulting MetTag CROP.seq LARRY-BC.ID library batch, hereafter referred to as MetTag-LARRY-GFP, was comparable to the original LARRY library (**Fig.S1B-D**). Next, we generated six unique isogenic ID8 p53^-/-^ murine ovarian cancer pools with MetTag-LARRY-GFP lentiviral libraries harboring unique batch identifiers from BC.ID1 to BC.ID6 (**Fig.1A**) and verified that the six lines had comparable *in vitro* proliferation rates (**Fig.S1E & S1F**).

**Figure 1:**
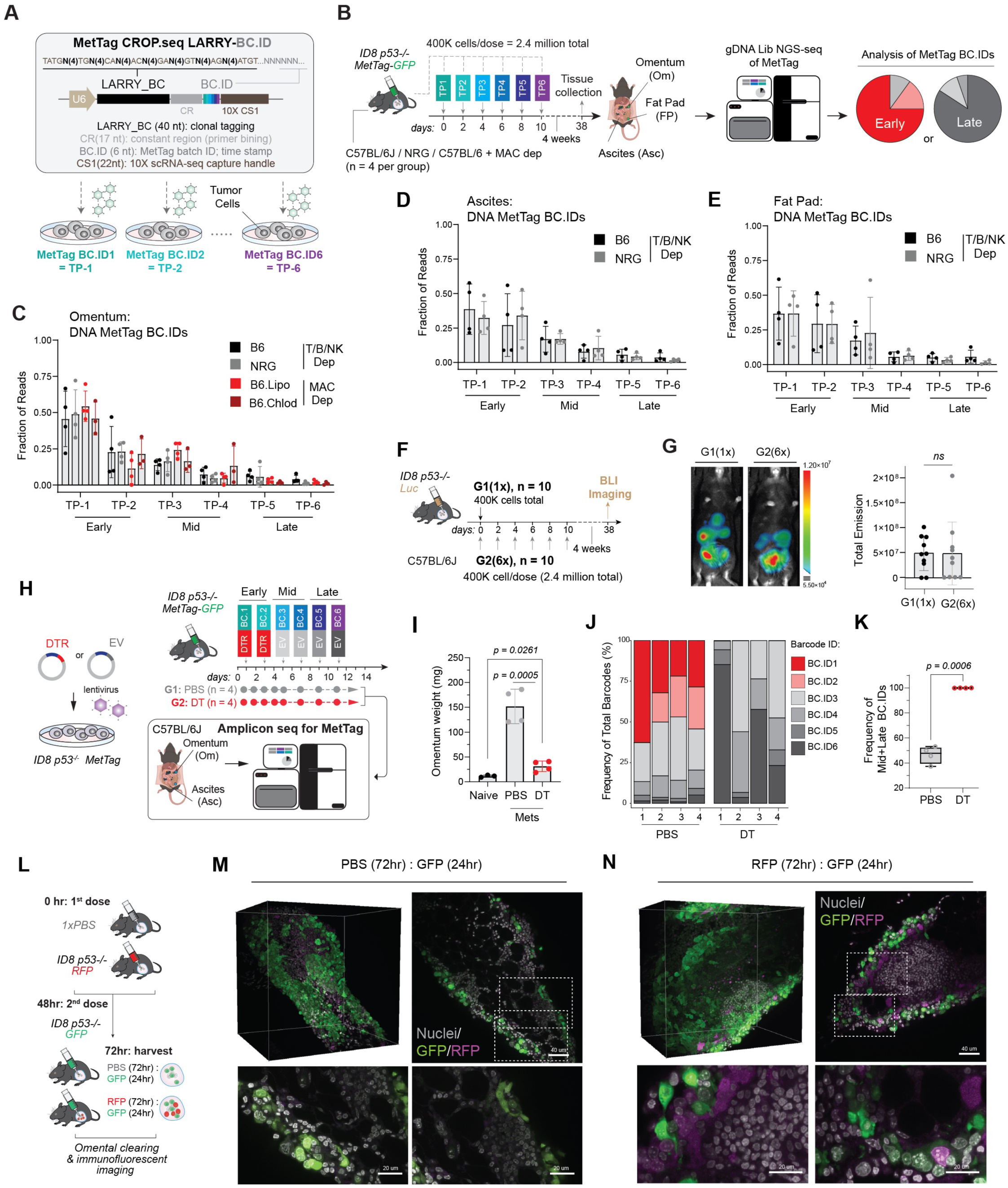
MetTag Captures Spatiotemporal Clonal Dissemination Patterns in Ovarian Cancer Metastasis. (**A**) Schematic depicting main elements of the MetTag CROP.seq LARRY-BC.ID, or MetTag-LARRY-GFP, barcoding construct. (**B**) Schematic of staggered injection and tissue collection timeline for clonal tracing using MetTag-LARRY-GFP barcoded ID8 p53^-/-^ ovarian cell line. Schematic was partially created with BioRender.com. (**C**) Genomic DNA (gDNA)-level barcode distributions in the metastatic omentum graphed as frequency under WT (n = 4), NRG host (n = 4), and MAC depletion conditions (control; n = 4, chlodrosome; n = 3). (**D-E**) gDNA-level barcode distributions in ascites and pelvic fat pads under WT (n = 4) and NRG hosts (n = 4). In (C-E) panels, BC.ID1 = TP-1, BC.ID2 = TP-2, BC.ID3 = TP-3, BC.ID4 = TP-4, BC.ID5 = TP-5, BC.ID6 = TP-6 for clarity. (**F**) Injection schema and timeline for comparing one-time (400,000 total, n = 10) cell dose and fractionated (2,400,000 total, n = 10) cell injections. (**G**) Representative BLI images at endpoint (left) and bar graph depicting total flux emission in each group at experimental endpoint. (**H**) Timeline schematic depicting *in vivo* DT-mediated ablation of DTR-expressing early DTCs, followed by NGS analysis of BC.ID (timepoint) distributions in metastatic omenta 14 days post initial cancer cell injection (n = 4 mice per group). (**I**) Quantification of omental weights at the 2-week experimental endpoint. Data was analyzed using a two-tailed Student’s t test and graphed as mean with standard deviation (SD). (**J**) NGS-derived BC.ID distributions depicted as frequency in DT and PBS-treated groups. (**K**) Contribution of Mid and Late BC.IDs in PBS and DT-treated groups. Data was compared using an unpaired Student’s t test with Welch correction and graphed as mean with SD. (**L**) Timeline schematic and groups used for early timepoint whole tissue omental imaging experiment. (**M-N**) Representative whole-tissue images of cleared omental tissues harvested 24 hours post GFP+ ID8 p53^-/-^ cell injection taken at 38x magnification. Images were taken in three representative locations per omenta (n = 4 mice total), without (M) or with (N) prior RFP+ ID8 p53^-/-^ cell preconditioning. 3D data was visualized using Napari software.

To model continuous dissemination *in vivo*, each individual MetTag-LARRY-GFP line was injected intraperitoneally (i.p.) in a fractionated manner at low doses (400,000 cells/dose, every 48 hours) across six different time points to model early/pioneer, mid, and late waves of DTCs (**Fig.1B**). Four weeks after the final DTC exposure (∼6 weeks after initial), ascitic fluid (Asc), omenta (Om), and peritoneal pelvic fat pads (FPs) were extracted. This experimental scheme was performed in several mouse models, including wild-type C57BL/6J mice, immunocompromised NOD.Cg-Rag1^tm1Mom^IL2rg^tm1Wjl^/SzJ (NRG) mice (lacking functional T, B, and NK cells), and under conditions of macrophage (MAC) depletion via chlodrosome administration. Temporally spaced DTC exposure resulted in robust metastasis formation in fully immunocompetent mice, evidenced by ascites accumulation and positive GFP signal in the omentum and pelvic adipose tissue (**Fig.S1G top**). Quantification of overall metastasis burden in a separate cohort of ID8 p53^-/-^ Luciferase cells injected into NRG mice yielded reduced omental burden compared to immunocompetent female C57BL/6 (B6) mice (**Fig.S1G & S1H**). Moreover, chlodrosome-mediated macrophage depletion in B6 mice resulted in reduced GFP+ cancer cell burden in ascites under the fractionated injection schema (**Fig.S1I & S1J**), suggesting a potential pro-metastatic role of both macrophages and lymphoid (T, B, and NK) cells in ovarian cancer.

We used MetTag-LARRY-GFP amplicon DNA sequencing to quantify the relative abundance of BC.IDs in final overt metastases: early timepoints (*BC.ID1=TP-1, BC.ID2=TP-2*), mid (*BC.ID3*=*TP-3, BC.ID4=TP-4*), and late (*BC.ID5=TP-5, BC.ID6=TP-6*) arrivers (**Fig.1B**, pie chart). Here, the term “late DTCs” refers to DTCs introduced at later timepoints into a system that has been preconditioned by prior waves of “mid” and “early” DTCs, and importantly, are not interchangeable with the clinical terms for early and late dissemination. Interestingly, in overt metastases collected from three distinct peritoneal niches (omentum, ascites, and fat pad), we observed a consistent pattern of early DTC enrichment, suggesting a relative dominance of early arrivers in shaping metastasis outcomes. We observed a nearly identical BC.ID distribution across all three anatomical locations in both wild type and NRG mice (**Fig.1C-1E**), as well as macrophage-depleted host in omenta (**Fig.1C**), suggesting a negligible role of niche-specific tumor immune microenvironment (TIME) in the overall dominance of early arriving DTCs, despite their apparent role in modulating overall metastasis burden.

### Early Arriving DTCs Are Sufficient to Drive Omental Colonization

To examine if later arriving cancer cells are dispensable for overall metastasis burden, we evaluated tumor burden in mice with a single injection of tumor cells (TP-1 only) compared to mice with fractionated dosing (TP-1-6) as described in Fig.1B in female B6 mice with luciferase-tagged ID8 p53^-/-^ cells. At the experimental endpoint of four weeks post final cancer cell injection, tumor burden levels were comparable (**Fig.1F & 1G**), as measured by *in vivo* bioluminescent imaging (BLI). This data indicates that early DTCs are not dependent on later DTCs for metastatic colonization and outgrowth. Next, we specifically depleted early DTCs via diphtheria toxin receptor (DTR)-mediated internalization of diphtheria toxin (DT). Upon validating DTR expression in MetTag-LARRY-GFP BC.ID1 and BC.ID2 lines and confirming DT sensitivity *in vitro* (**Fig.S1K & 1L**), we conducted a similar staggered injection experiment as described in Fig.1B (400,000 cancer cells/dose) with the addition of either PBS or DT treatment and collected omenta at an early week 2 timepoint (**Fig.1H**). Depletion of early DTCs via DT treatment significantly reduced omental metastatic burden (**Fig.1I**), with proportional dominance of mid and late BC.IDs (**Fig.1J & 1K**). Importantly, given that this BC.ID compositional shift did not result in aggressive outgrowth of later-arriving DTCs (**Fig.1I**), these results indicate that early DTCs are functionally essential and largely contribute to full metastatic outgrowth. Furthermore, the sequential seeding experimental scheme prompted us to ask whether observed early DTC dominance reflects an advantage of seeding kinetics rather than mechanisms residing in DTC-intrinsic adaptation and/or PSMN co-evolution. To determine whether 24 hours is sufficient for omental colonization and whether prior seeding of DTCs constrains subsequent seeding and colonization, we performed a short-term (72 hours) *in vivo* sequential injection experiment using RFP-labelled cells followed by GFP-labelled ID8 p53^-/-^ cells (**Fig.1L**). Whole-tissue immunofluorescence light-sheet microscopy performed 24 hours after GFP-cell injection showed that RFP+ cells (TP-2) were able to colonize the omentum at similarly detectable levels regardless of prior GFP+ cell seeding (**Fig.1M & 1N**). These data indicate that prior seeding by leading DTCs (GFP+) does not appreciably compromise the ability of subsequently arriving DTCs to colonize the omentum over the first 72 hours.

### Clonal Analysis Reveals Shared Evolutionary Path Towards an Inflammatory Signature

To gain a better understanding of the transcriptional programs necessary for early DTCs to colonize and subsequently assess their clonal expansion status, we performed CITE-seq (scRNA-seq with HTO sample multiplexing) coupled with direct capture of U6-driven MetTag-LARRY-GFP transcripts via capture sequence 1 (CS1) (**Fig.2A**). Sequential injections were performed in the same manner as in Fig.1B with a reverse barcode injection schema to ensure no unintended bias towards certain barcodes gained over the cell selection period. Single cell analysis of MetTag-LARRY-GFP BC.IDs from metastasis-bearing tissues revealed that early DTCs (*TP1-2*) were enriched at endpoint (**Fig.2A & 2B**). The prominent trend towards early DTC dominance was consistent with gDNA sequencing data in Fig.1C and 1D. We next asked whether the dominance of early DTCs in overt metastases represents a widespread niche-adaptive and PSMN conditioning feature of DTCs. We applied three MetTag-LARRY-GFP BC.IDs in a triple negative breast cancer (TNBC) mouse model of lung metastasis (Lung.Met): BC.ID1 representing early TP-1 and TP2, BC.ID2 representing mid TP-3 and TP4, and BC.ID6 representing the late TP-5 and TP-6. In this model, we identified almost exclusively early TP DTCs dominance (91.9%) in overt lung metastases (**Fig.2C**). Collectively, these data suggest that early-arriving DTCs preferentially dominate endpoint metastases in both hematogenous (TNBC) and transcoelomic (ovarian) metastases.

**Figure 2:**
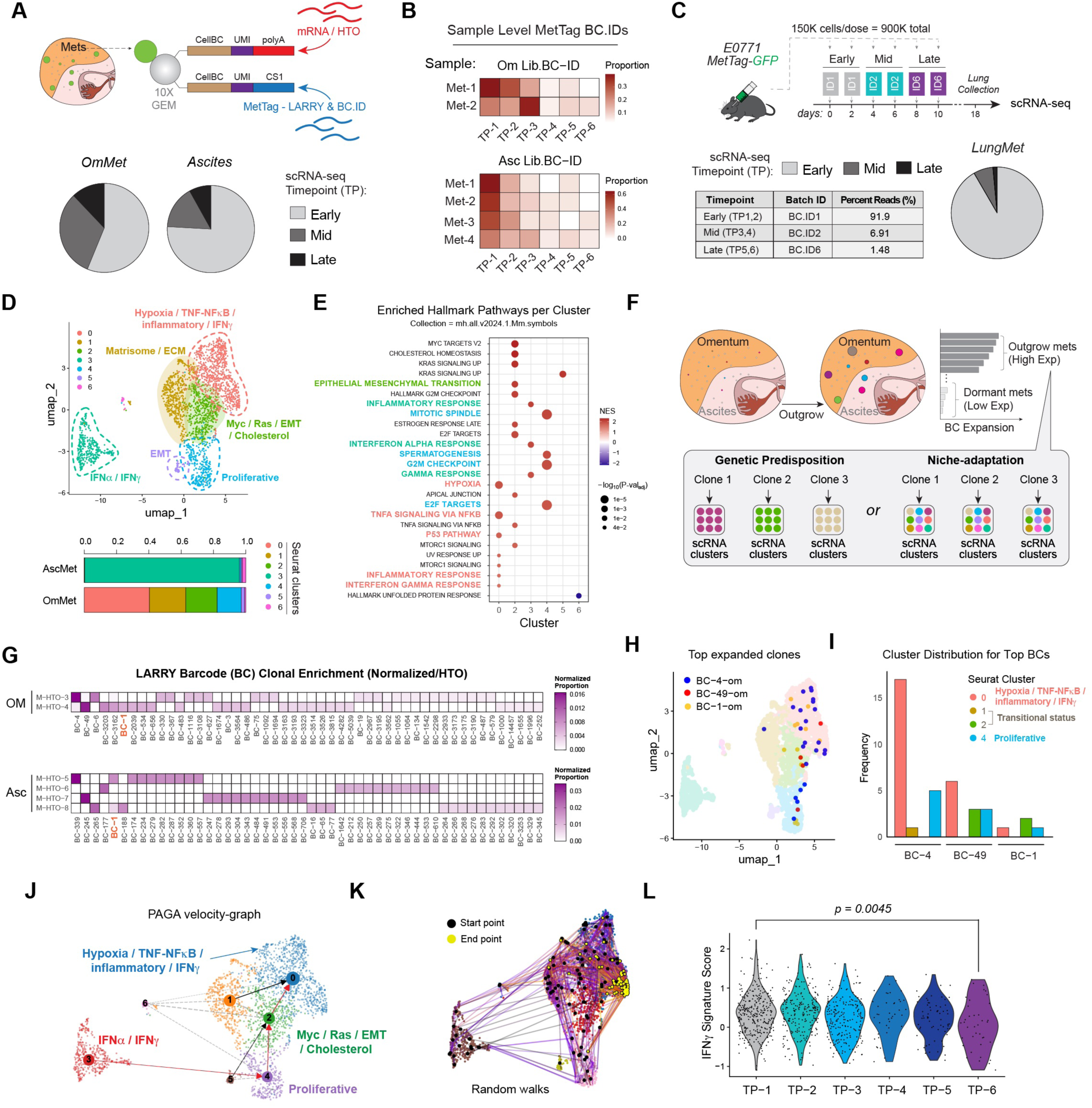
Clonal Analysis Reveals Shared Evolutionary Path Towards Inflammatory Signature. (**A**) Schematic of scRNA-seq-based MetTag-LARRY-GFP and transcriptome detection method where LARRY barcodes are readily captured by Chromium 10x Feature Barcoding (top). Pie charts of RNA-level BC.ID frequencies in each location (bottom) grouped as early (BC.ID5, BC.ID6), mid (BC.ID4, BC.ID3), and late (BC.ID2, BC.ID1). This experiment was carried out with a reversed BC.ID injection order as compared to Figure 1. OmMet; HTO n’s = 2, Ascites; HTO n’s = 4. (**B**) Heatmap of barcode frequencies, per individual sample/HTO: BC.ID6 = TP-1, BC.ID5 = TP-2, BC.ID4 = TP-3, BC.ID3 = TP-4, BC.ID2 = TP-5, BC.ID1 = TP-6. (**C**) Timeline schematic (top) and MetTag-LARRY-GFP BC.ID distribution (bottom) in independent TNBC lung metastasis model depicting enrichment of scRNA-seq derived early TP DTCs in metastatic lungs (HTO n’s = 3) at endpoint. (**D**) Seurat cluster UMAP of reclustered cancer cells with transcriptional program labels obtained from integrated OmMet and Ascites samples (top). Stacked bar chart depicting the frequency of Seurat subclusters within each metastatic location (bottom). (**E**) Dot plot depicting Hallmark pathways across all subclusters used to define each cluster subset gene program in (D). (**F**) Schematic representation of the possible outcomes of LARRY clonal analysis based on transcriptome patterns of enriched clones. (**G**) Heatmap of normalized clonal LARRY barcode counts across the two anatomical locations and individual mice (HTOs). (**H**) UMAP overlay of top 3 expanded LARRY clones in the omentum. (**I**) Bar chart depicting the relative subcluster proportions present in each of the top 3 barcode clones. (**J**) scVelo-generated PAGA velocity graph indicating trajectory arrows and inferred cluster lineage. (**K**) scVelo and CellRank-based simulated path from start to endpoint clusters. (**L**) Violin plot depicting RNA-level contribution of each timepoint to the IFNγ signature. IFNγ signature was defined using canonical mouse genes, followed by directly applying the wilcox.test() function in Seurat to compare TP-1 and TP-6 expression.

As most DTCs do not survive and only rare clones successfully adapt to distant niches^19^, we examined whether tumor cells disseminated over time evolve along diverse trajectories or converge on a common path characterized by a limited set of adaptive transcriptome features. We first annotated cell clusters with higher levels of keratins and absence of CD45 transcript expression (*Krt18*+, *Krt8*+, *Ptprc*-) as the tumor cell-enriched cluster on the UMAP (**Fig.S2A & S2B**). Tumor cell cluster states were determined through a combination of cluster marker genes and pathway-level analysis using the escape R package^20^ (**Fig.S2C – S2E**). Ascites-derived cancer cells were strongly enriched for interferon alpha (IFNα) and gamma (IFNγ) signaling (cluster 3), as opposed to four distinct heterogeneous transcriptional states in OmMet including proliferative, epithelial-mesenchymal transitioning (EMT), matrisome, and a hypoxic, inflammatory subcluster (**Fig.2D & 2E**).

Cellular barcoding approaches have been widely used to reveal clonal dynamics driven by both genetic predisposition and epigenetic changes in response to environmental cues^21^. As even isogenic cell lines consist of preexisting epigenetically diverse cell populations^22^, we asked whether the most expanded clones would 1) cluster together based on transcriptional status, indicating strong pre-determined cell-autonomous programs driving metastasis, or 2) disperse into heterogeneous transcriptional states (clusters) reflective of diverse adaptations to different metastatic niche cues (**Fig.2F**). First, we observed minimal overlap between dominant outgrowing clones across each mouse (HTO sample) in both anatomical locations (**Fig.2G**). While one clone, BC-1, was shared across ascites and omenta, most of the highly expanded clones are distinct between niches. Although these results do not exclude genetic contributions, they suggest that successful clonal outgrowth requires unique adaptations reflective of the specific microenvironment.

Given that omental metastases showed diverse single-cell transcriptional states (**Fig. 2D-E**), we projected clonal identity of the top three expanded clones from omenta onto the cancer cell UMAP and examined similarities between clones. Importantly, the top three identified clones, BC-1, BC-4, and BC-49, were not significantly overrepresented in the six MetTag-LARRY-GFP lines before injection (**Fig.S2F**). At both the cluster and pathway levels *in vivo*, dominant clones were enriched for several defined signatures (**Fig.2H-2I; S2G**). Apart from the proliferative subcluster 4, expanded clones consistently occupied the hypoxic, inflammatory subcluster 0 (**Fig.2I**), suggesting diverse, yet directed evolutionary trajectories within the same clonal population. To understand the relationships among different subclusters, we applied RNA velocity and pseudotime analysis using scVelo and CellRank^23^. This approach revealed a clear predicted trajectory originating from the IFN signaling enriched ascites compartment subcluster 3, leading to proliferative status, then further converging to subcluster 0, a terminal inflammatory state in the omentum (**Fig.2J & 2K; Fig.S2H**). Notably, IFNγ, but not the IFNα signature, was significantly enriched in early TP-1 DTCs (**Fig.2L; S2I**).

Similarly to the ovarian cancer dataset, we observed a heterogeneous clonal distribution pattern between each mouse in the TNBC lung metastasis MetTag-LARRY-GFP scRNA-seq dataset, suggesting non-genetic, transcriptomic drivers of successful colonization (**Fig.S2J & S2K**). To examine the generality of the enriched IFN signature in metastases, we performed a separate scRNA-seq experiment comparing TNBC Lung.Met and E0771 cells *in vitro* (**Fig.S3A**). Compared to cultured cells, *in vivo* Lung.Met tumor expressed higher IFNγ and IFNα signatures (**Fig.S3B-S3E**). Collectively, the ovarian and lung metastasis data suggests that transcriptional adaptation towards IFN-signaling *in vivo* is a common trait of successfully colonized DTCs. In ovarian metastasis specifically, successful metastatic clones in the omentum occupy heterogeneous transcriptional states. scRNA-seq and trajectory analyses further suggests that these omental states may arise from an IFN-signaling-rich ascites population that precedes later niche-specific adaptations.

### *In vivo* CRISPR screen identifies functional essentiality of IFNγ signaling in omental colonization

We next asked whether transcriptional adaptation of cancer cells to the ascites environment is essential for subsequent intraperitoneal seeding and successful omental metastasis. We hypothesized that the ascites gene signature adopted by early DTCs – characterized by limited nutrient availability and low attachment – is (1) induced by initial detachment from a primary tumor and (2) necessary for subsequent peritoneal metastasis. To test this hypothesis, we performed differential gene expression (DEG) analysis and over-representation pathway analysis (ORA) with the clusterProfiler R package^24^ to compare gene expression in metastatic tumor cells residing in the two distinct niche compartments (Asc vs Om). Compared to OmMet, DEGs of AscMet consisted of hemoglobin genes (*Hbb-bs, Hba-a1, Hba-a2, Hbb-bt*) and interferon-stimulated genes (*Ifi27l2a* and *Irf7*). In contrast, OmMets were enriched in immediate-early response genes (*Atf3, Egr1, Fos, Fosb, Jun*)^25,26^, adaptive immune immunoglobulin components (*Igkc, Igha, Jchain*)^27^, and angiogenesis (*Vegfa*) (**Fig.3A**). Next, IFNγ/IFNα module scoring and pathway analysis confirmed a dominant interferon-rich inflammatory signature in AscMet, as opposed to more heterogeneous programs in OmMet (**Fig.3B; Fig.S3F-H**).

**Figure 3:**
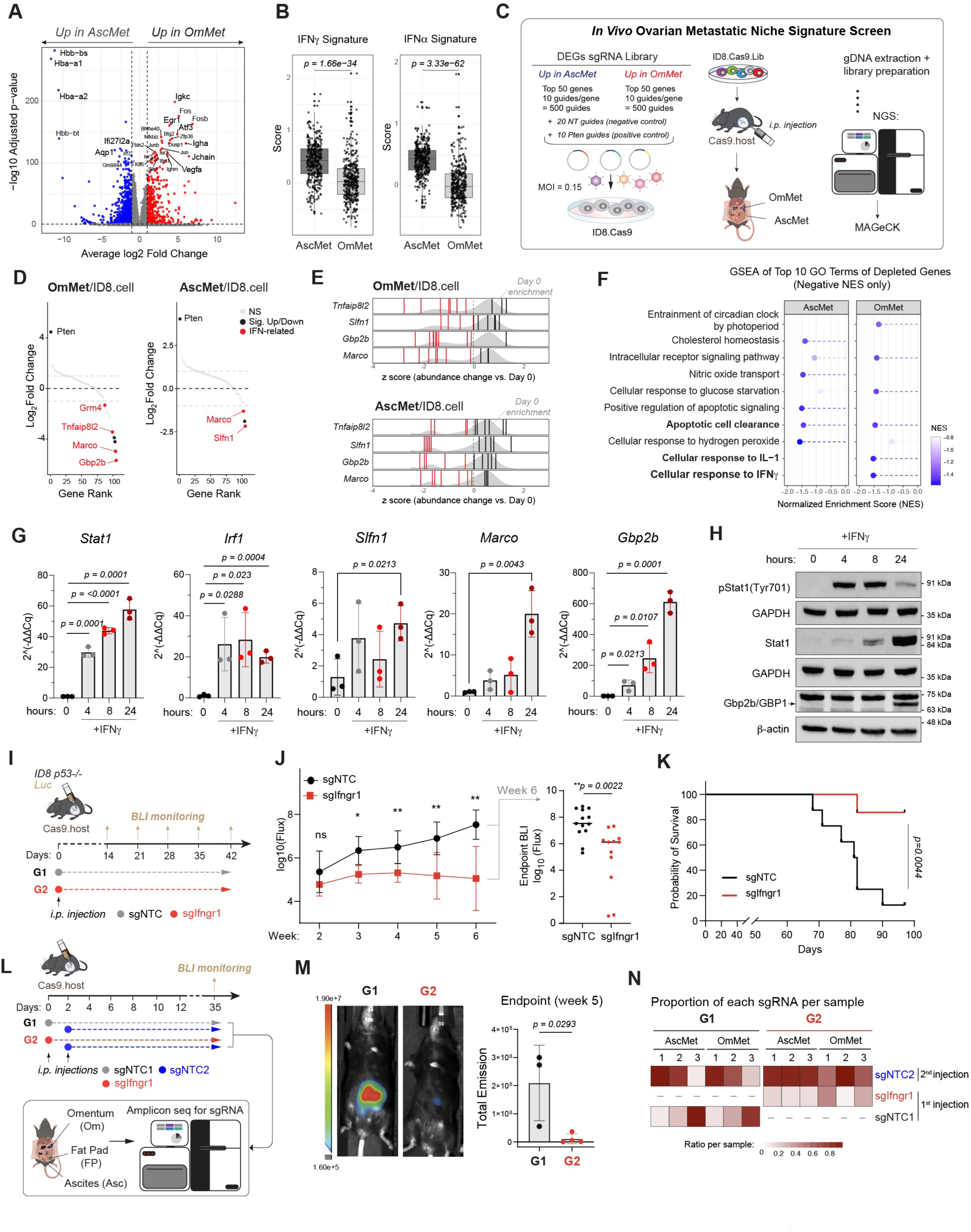
Early DTC Activation of IFNγ Response is Necessary for Successful Omental Colonization. (**A**) DEG-based volcano plot of gene expression differences between OmMet and AscMet, generated using the FindMarkers() function where threshold for significance was defined at p <0.05 and log2FC < -1.0 for downregulation or >1.0 for upregulation. (**B**) Boxplots comparing IFN signature modules between AscMet and OmMet. Custom IFN signatures were defined in RStudio, followed by wilcox.test() in Seurat to compare module scores between AscMet and OmMet. (**C**) Schematic describing *in vivo* ovarian metastatic niche signature screen. Partially created with BioRender.com. (**D**) RankView plots for gene-level log2 FC changes in OmMet (n=5, left) and AscMet (n=4, right) samples relative to Day 0 cancer cells. Red dots signify depleted interferon-related genes. (**E**) Distribution of 10 individual sgRNAs targeting depleted interferon-related genes. Gray histogram represents global distribution of guides at baseline (day 0) and red bars indicate depleted guides with z-score < 0. (**F**) Lollipop plot of top 10 depleted pathways associated with depleted genes in (E). CRISPR screen related data (3D-3F) was analyzed using MAGeCK and MAGeCKFlute. (**G**) IFNγ treatment time course qPCR fold change (2^(-ΔΔCq)) plots of select CRISPR hits and positive controls. Each data point represents a biological replicate (n = 3), and each biological replicate is computed as an average of individual technical replicates. Data was analyzed using a two-tailed Student’s t test and plotted as mean with SD in Prism. (**H**) IFNγ treatment time course western blot probing for Stat1/pStat1 and GBP1. (**I**) Timeline schematic of experimental metastasis study comparing sgNTC and sgIfngr1.g1 groups, each with one time injection of 400,000 ID8 p53^-/-^ Luciferase cells. Two independent groups pooled; cohort 1 sgNTC; n = 8, sgIfngr1; n = 7; cohort 2 sgNTC; n = 5, sgIfngr1; n = 5. (**J**) Time course log10-transformed total emission/flux values (left). log10-transformed flux plot for week six with p value (right). Data was analyzed using a two-tailed Student’s t test and plotted as median with individual values in Prism. (**K**) Kaplan Meier survival curve for cohort 1 sgNTC (n = 8) and sgIfngr1 (n = 7). Survival data was computed using a standard log-rank test. (**L**) Timeline schematic for staggered sgIfngr1 (n = 4) or sgNTC1 (n = 3) followed by sgNTC2-cell injection experiment with BLI monitoring. (**M**) Representative endpoint BLI images of metastasis bearing mice in each group (left) and endpoint total BLI flux quantification comparing sgIfngr1 and sgNTC first injection (right). Data was analyzed using a two-tailed Student’s t test and plotted as mean with SD in Prism. (**N**) gDNA-level guide NGS-derived heatmap depicting relative enrichment of 1^st^ DTC wave - either sgNTC1 (G1) or sgIfngr1 (G2) – compared to the 2^nd^ DTC wave, represented by sgNTC2.

To assess functional significance of AscMet and OmMet DEGs, we designed a custom *in vivo* CRISPR/Cas9 screen targeting 100 DEGs (**Fig.3C**). SpCas9 was introduced into ID8 p53^-/-^ cancer cells and the functionality of the CRISPR system was validated *in vitro* (**Fig.S3I & S3J**). 2.4 million cells (∼2,000X representation of sgRNA library) were i.p. injected to a whole-body Cas9-expressing host (**Fig.3C**). This host has central tolerance to Cas9, decreasing immunogenicity towards Cas9-expressing tumor cells^28^. Six weeks post-injection, metastasis-bearing omenta and ascites were harvested for genomic DNA (gDNA) extraction and NGS sequencing, followed by MAGeCK^29^ analysis (**Fig.3C**). The positive controls for the screen, *Pten* gRNAs, were significantly enriched in metastases derived from both anatomical locations (**Fig.3D**). Compared to the initial injected cell pool sgRNA distribution, sgRNAs targeting eight genes in OmMet and ten genes in AscMet were significantly depleted (*p < 0.05*) (**Fig.3D & 3E**). The top depleted (dropout) genes – *CD300lb*, *Ms4a8a*, *Marco*, *Gbp2b* (OmMet), and *Slfn1, Marco* (AscMet) – were upregulated DEGs associated with AscMet, suggesting a necessity of initial transcriptome adaptation to the ascites niche - the first niche DTCs encounter - for subsequent metastasis. Furthermore, GSEA analysis of dropout genes using the GO Biological Function (GOBP) gene sets revealed a shared depletion of apoptotic cell clearance processes in both Asc and Om compartments and drop out of programs related to IFNγ in omentum (**Fig.3F**).

To experimentally validate whether IFNγ is required for the transcriptional upregulation of CRISPR screen hits in cancer cells, we treated ID8 p53^-/-^ cells *in vitro* with recombinant IFNγ. We tested the inducibility of *Marco*, *Gbp2b*, and *Slfn1* by IFNγ via RT-qPCR, where *Stat1* and *Irf1* transcription factors served as positive controls for IFNγ response^30^ (**Fig.3G**). By 24 hours of treatment, all three genes were significantly induced by IFNγ (**Fig.3G**). The protein level of Gbp2b, an interferon-inducible GTPase, was also significantly induced by IFNγ (**Fig.3H**). Collectively, despite the transcriptional heterogeneity revealed by scRNA-seq, consistent enrichment of apoptotic pathway and the sequential activation of an IFNγ response (first in ascites, then required for omental metastasis) suggest two functionally essential adaptive mechanisms employed by DTCs to survive *in vivo* clonal selection.

### Loss of IFNγ-response in DTC significantly decreases peritoneal metastases and extends overall survival

As most of the top hits from the CRISPR screen were IFNγ-inducible, we reasoned that IFNγ may serve as a primary regulator that exerts broader pro-metastatic adaptive cellular programs in DTCs. Thus, instead of focusing on a single downstream gene target, we sought to disable the entire cascade using an IFNγ receptor 1 (*Ifngr1*) knockout. We first confirmed robust expression of *Ifngr1* in both AscMet and OmMet *in vivo* (**Fig.S3K**). Next, we genetically knocked out *Ifngr1* in ID8 p53^-/-^ cells using three individual guides (sg#1, sg#4, and sg#6). Successful knockout of *Ifngr1* was validated via flow cytometry (**Fig.S3L**) and ablation of the IFNγ transcriptional downstream targets was validated via western blot probing for Stat1 and Gbp2b for sg#1 (**Fig.S3M**). We next i.p. injected ID8.Luciferase-sgNTC or ID8.Luciferase-sgIfngr1(sg#1) to Cas9-expressing mice. Tumor burden was monitored by BLI intensity throughout the 6-week experimental window (**Fig.3I**). sgIfngr1 groups showed a significant reduction of overall metastasis burden compared with sgNTC group (**Fig. 3J; Fig.S3N**). In contrast to sgNTC group, ∼80% of mice in sgIfngr1 groups remained alive at the end of the survival study (p = 0.004) (**Fig.3K**).

Above results demonstrated that DTCs’ intrinsic IFNγ signaling is essential for the overall metastatic outcome. Given that early DTCs (TP-1) showed significantly higher IFNγ signaling compared with later DTCs (**Fig.2L**), we next investigated whether the IFNγ response in early DTCs influences the success of subsequent waves of DTC colonization. We specifically perturbed IFNγ response (sgIfngr1) in the first batch of early DTCs (G2), but not the immediate second wave of sgNTC2 DTCs (**Fig.3L**). Interestingly, loss of Ifngr1 in the first batch of DTCs significantly reduced overall metastasis burden at experiment endpoint (**Fig.3M**). In the residual metastases in the sgIfngr1 group (G2), control (sgNTC2) cells outcompete the sgIfngr1 cells, as shown with sgRNA amplicon sequencing (**Fig.3N**). Taken together, these results demonstrated that, although a robust IFNγ response in pioneer DTCs is associated with their relative enrichment over later DTCs, this early response is paradoxically required for overall metastasis success (**Fig.3L-N**).

### IFNγ response mediates resistance to anoikis by activating pro-survival programs

We next sought to examine how IFNγ signaling promotes metastasis. First, we observed that *Ifngr1* knockout did not alter cancer cell fitness under standard adherent culture conditions in the presence or absence of IFNγ treatment, evidenced by both CellTiter-Glo luminescent cell viability assay (**Fig.4A**, left) and colony count formation (**Fig.S4A**). Interestingly, IFNγ treatment had a protective effect and increased cell viability in control (sgNTC) cells under ultra-low attachment (ULA) conditions. This protective effect was ablated in the *sgIfngr1* groups (**Fig.4A**).

**Figure 4:**
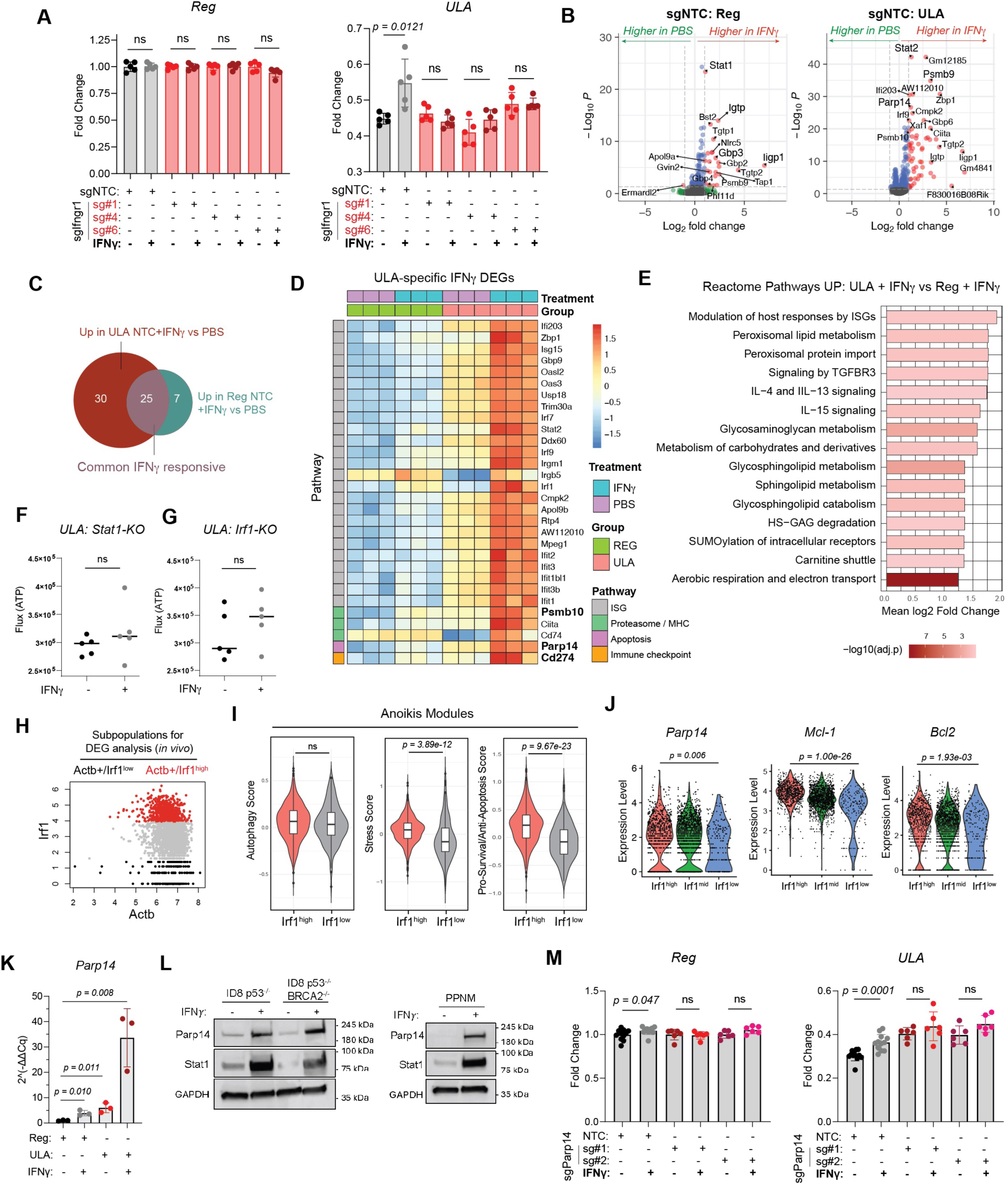
IFNγ Response Mediates Resistance to Anoikis Through Activation of Anti-apoptotic Programs. (**A**) Cell-Titer Glo (CTG) assay luminescence assay graphed as fold change for regular culture conditions (left) and ultra-low attachment (ULA) conditions (right), with and without IFNγ treatment, comparing viability of sgNTC and sgIfngr1 (sg#1, sg#4, and sg#6) lines. Fold change for each biological replicate (n = 5) was determined by normalizing each cell line’s CTG value to the regular attachment, without IFNγ treatment, baseline. CTG assay luminescence data was analyzed using a two-tailed Student’s t test and graphed as mean with SD. (**B**) Volcano plots generated from bulk RNA-seq depicting top DEGs between sgNTC (wild type) cells under regular culture conditions with (n = 3) or without (n = 3) IFNγ treatment (left) and under ULA conditions with (n = 3) and without (n = 3) IFNγ treatment (right). Bulk RNA-seq data was analyzed using DESeq2 and visualized as variance-stabilized counts. (**C**) Venn diagram showing 25 overlapping and 30 unique upregulated genes in sgNTC cells treated with IFNγ under ULA conditions. (**D**) Heatmap of vst-normalized gene counts for ULA-specific upregulated genes, ordered by treatment (PBS, IFNγ) and group (Reg, ULA). (**E**) Bar plot depicting top 15 Reactome pathways upregulated under ULA conditions treated with IFNγ compared to regular with IFNγ. CTG flux dot plots for Stat1-KO (**F**) and Irf1-KO (**G**) cells under ultra-low attachment, with (n = 5) or without (n = 5) IFNγ treatment. Data in (F) and (G) was graphed as median with individual values and analyzed using a two-tailed Student’s t test. (**H**) *In vivo* scRNA-seq based gating strategy used to identify Irf1^high^ and Irf1^low^ cells for subsequent module scoring. (**I**) Violin plots comparing autophagy, stress, and anti-apoptosis module expression in Irf1^high^ and Irf1^low^ cells. (**J**) Violin plots comparing expression level of Parp14 (left), Mcl-1 (middle), and Bcl2 (right) between Irf1^high^ and Irf1^low^ cells. scRNA-seq based transcript expression levels depicted in (I) and (J) were compared by applying the wilcox.test() function in Seurat. (**K**) RT-qPCR derived Parp14 transcript levels in ID8 p53^-/-^ cells under adherent (Reg) and detached (ULA) culture conditions and IFNγ treatment, n = 3 per group. Fold change (2^(-ΔΔCq)) data was analyzed using a two-tailed Student’s t test and graphed as mean with SD. (**L**) Protein level validation of Parp14 inducibility in detached (ULA) culture in three murine cell lines. (**M**) Cell-Titer Glo (CTG) assay luminescence assay graphed as fold change for regular culture conditions (left) and ULA conditions (right), with and without IFNγ treatment, comparing viability of sgNTC and sgParp14 (sg#1 and sg#2) lines. Fold change for each biological replicate (n = 6) was determined by normalizing each cell line’s CTG value to the regular attachment, without IFNγ treatment, baseline. CTG assay luminescence data was analyzed using a two-tailed Student’s t test and graphed as mean with SD.

To investigate the underlying mechanism, we performed bulk RNA-seq (**Fig.S4B**) with wild type (sgNTC) and single clone knockout line (sgIfngr1, sg#1). First, principal component analysis (PCA) and MA dispersion plot showed that ULA culture alone induced significant transcriptomic shifts in both sgNTC and sgIfngr1 cells (**Fig.S4C & S4D**). Wild type cells that survived under ULA conditions exhibited an overall reduced S/G2 signature (**Fig.S4E**). Next, we compared the transcriptomic changes in response to IFNγ. PCA of all groups revealed variance between wild type cells treated with PBS and IFNγ. This IFNγ induced variance was ablated in the sgIfngr1 knockout line as expected, independent of culture conditions (**Fig.S4C**). We analyzed DEGs between cells with or without IFNγ treatment under both regular and ULA conditions (**Fig.4B**). Interestingly, despite lower proliferation, cells grown under ULA condition exhibited transcriptionally diverse responses (**Fig.4B; Fig.S4F**) and were overall more responsive to IFNγ than under regular condition, rendering more DEGs detected (**Fig.4C**). Besides 25 common IFNγ responsive genes, we discovered 30 IFNγ responsive genes uniquely up-regulated only under ULA condition (**Fig.4C**). We next grouped the unique 30 upregulated genes into broad classes. As expected, most genes represented classical interferon stimulated genes (ISGs) with known roles in viral defense and antigen presentation (**Fig.4D**). In addition, we discovered several non-canonical ISGs, such as the immunoproteasome 20S subunit beta 10 (*Psmb10*), poly(ADP-ribose) polymerase family member 14 (*Parp14*), as well as PD-L1 (*Cd274*) (**Fig.4D**). Pathway level comparison between IFNγ-treated cells under ULA versus adherent (Reg) conditions revealed Reactome pathways related to metabolic rewiring and glycosaminoglycan remodeling (**Fig.4E**). Importantly, Psmb10, Parp14, and PD-L1 all have reported context dependent, cell autonomous pro-survival functions^31–34^. Supporting the functional role of downstream responsive genes, knockout of IFNγ transcription factors *Stat1* and *Irf1* phenocopied Ifngr1 knockout, leading to loss of IFNγ-induced survival phenotype in suspended cultures (**Fig.4F & 4G; Fig.S4G**).

### Parp14 is a Critical IFNγ-induced Effector of DTC Fitness

To validate *in vivo* relevance of these programs, we analyzed above DEGs detected in bulk RNA-seq using our single cell RNA-seq data derived from ovarian metastases. *In vivo* AscMet exhibited higher levels of immunoproteasome genes (*Psmb10*, *Psmb9*) as well as poly(ADP-ribose) polymerase family members (*Parp12*, *Parp14*, and *Parp9*) compared with OmMet (**Fig.S4H**). Unexpectedly, *Cd274* was more highly expressed in OmMet (**Fig.S4H**). Next, we performed RNA-based gating and defined *Irf1*^high^ and *Irf1*^low^ cancer cells to represent IFNγ downstream responsive levels of each cell (**Fig.4H)**. We curated custom module scores that reflected common pathways cancer cells utilize to resist anoikis-mediated death: autophagy, stress, and pro-survival/anti-apoptosis^35^ (**Fig.4I**). *Irf1*^high^ cells most significantly upregulated the pro-survival/anti-apoptosis module (**Fig. 4I**, right panel). On an individual gene level, *Irf1*^high^ cells upregulated *Parp14, Mcl-1* and *Bcl2*, but not *Cd274* and *Psmb10* (**Fig.4J** & **Fig.S4I**).

Finally, we assessed the functional role of Parp14, a known inhibitor of apoptosis in multiple myeloma^33^. We confirmed robust IFNγ inducibility of Parp14 transcript and protein under ULA conditions *in vitro* (**Fig.4K & 4L**), followed by CRISPR-mediated knockout (**Fig.S4J**). Interestingly, while baseline Parp14 levels sensitized cancer cells to suspension-induced stress without IFNγ, partial <u>knockout abolished the pro-survival phenotype induced by IFNγ treatment</u> (**Fig.4M**), indicating that <u>IFNγ induction of</u> Parp14 is necessary to facilitate DTC survival.

### Early DTC niche pre-conditioning leads to an LPM and NK-rich microenvironment

Given that early pioneer DTCs possess a unique survival advantage through IFNγ signaling, we next explored if initial dissemination events remodel the metastatic niche at an early timepoint. We performed flow cytometry analysis on ascites harvested from the two-week early time point DT treatment experiment (**Fig.1H**). In the absence of early DTC depletion (PBS group), metastasis-bearing ascites exhibited significant shifts in both myeloid and lymphoid cell populations relative to the early-DTC-depleted (DT-treated) group. Within the myeloid compartment, depletion of early DTCs significantly reduced the frequency of large peritoneal macrophages (LPMs) in ascites compared with the PBS control group, whereas small peritoneal macrophages (SPMs) and neutrophils were largely unchanged (**Fig.5A**). Within the lymphoid compartment, NK cells were significantly reduced in the DT-treated group, whereas total T cells increased. The CD8/CD4 T cell ratio remained comparable between the two groups (**Fig.5B**).

**Figure 5:**
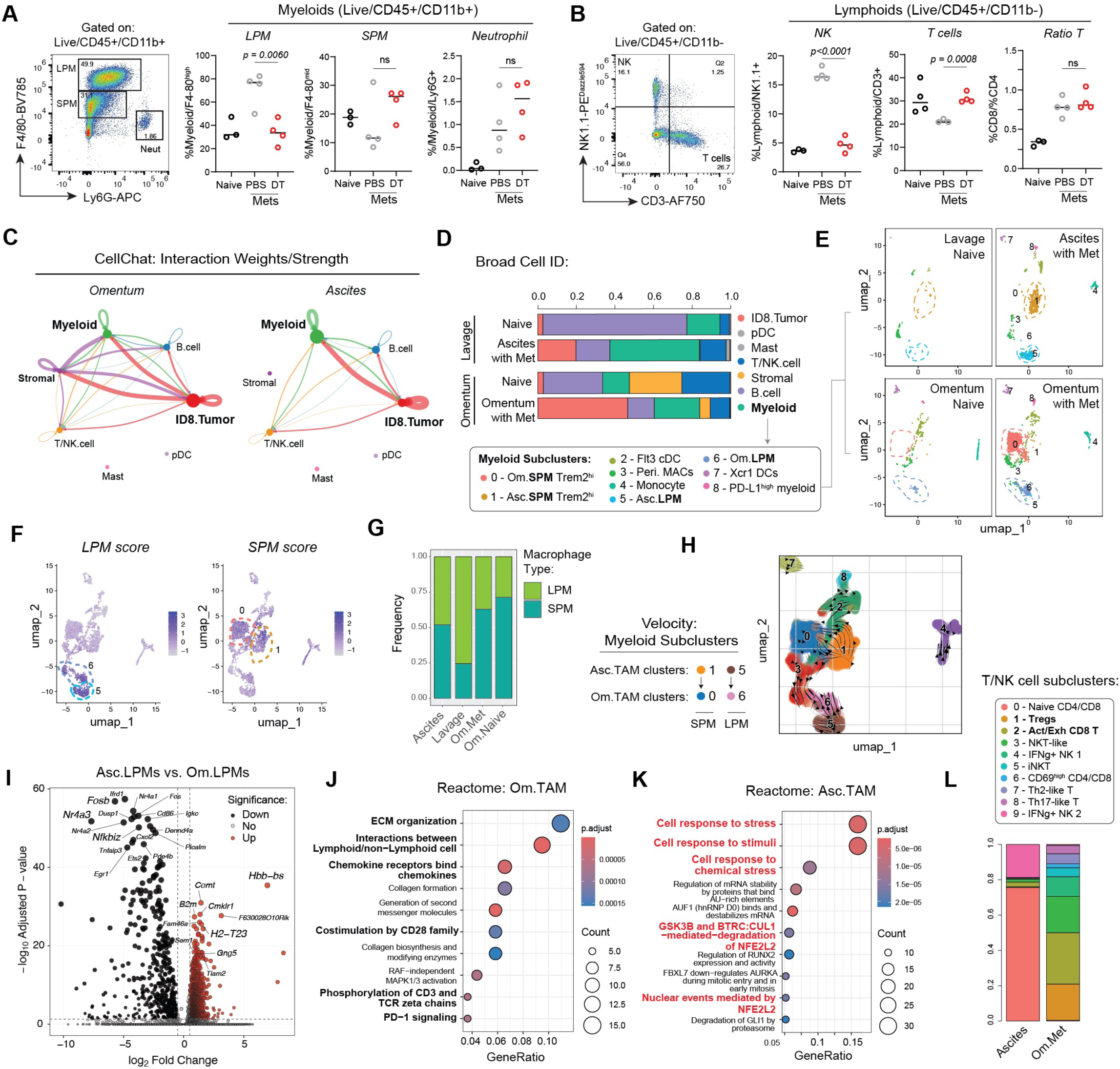
Pioneer DTC Colonization Shapes Niche-specific Macrophage States. Flow cytometry assessment of (**A**) myeloid and (**B**) lymphoid cell populations in naïve (n = 3), PBS-treated (n = 4), and DT treated (n = 4) B6 mice two weeks post initial cell injection. Timeline schematic for the 2-week DTR-mediated early DTC wave ablation experiment can be found in Figure 1H. Flow cytometry data was first analyzed in FlowJo, followed by two-tailed Student’s t test analysis in GraphPad Prism. Data was graphed as median with individual values. (**C**) CellChat-generated circos plots of interaction strengths between Broad cell ID categories in metastatic omenta (left) and ascites (right), derived from scRNA-seq. (**D**) Stacked bar chart of cell type proportions across the four experimental groups: naïve lavage (HTO n’s = 4), ascites (HTO n’s = 4), naïve omenta (n =4 pooled into one HTO), and metastatic omenta (HTO n’s = 2). (**E**) UMAPs of reclustered myeloid cells, split by experimental condition. (**F**) Feature plots depicting UMAP coordinates of cells with high LPM and SPM signature expression (darker purple). (**G**) Stacked barchart of relative LPM/SPM ratios in each experimental group. (**H**) UMAP of subclustered myeloid cells overlayed with scVelo RNA velocity arrows. Arrows are pointing from ascites TAMs (clusters 1 and 5) to omentum TAMs (clusters 0 and 6). (**I**) DEG volcano plot comparing gene expression differences between ascites LPMs and omentum LPMs. Significance of DEGs in Figure (I) was evaluated by applying the FindMarkers() and Wilcoxon test in Seurat to subsetted LPMs. Dot plots depicting ORA Reactome pathways enriched in ascites (**J**) and omentum-derived TAMs (**K**), subsetted as *Cd68*^high^/*Adgre1*^high^ macrophages. (**L**) Stacked bar chart of subsetted T/NK cells (Broad.cell_ID) comparing lymphoid cell type proportions in ascites and omentum.

Next, we assessed major interaction hubs *in vivo* at metastatic endpoint using CellChat^36^ analysis on major cell types derived from scRNA-seq Seurat clusters (**Fig.S5A & S5B**). Although there was a general increase in both the number and strength of cell-cell interactions between cells from naïve and metastasis-derived samples (**Fig.S5C**), tumor cells primarily interacted with myeloid cells in both ascites in omentum (**Fig.5C**). Stromal interactions were enriched specifically in the metastatic omentum (**Fig.5C**). This suggests that interaction between IFNγ-responsive tumor cells and macrophages may be a key axis of communication across niches. We found a proportional increase in myeloid cells in both metastatic sites compared to their naïve counterparts (**Fig.5D**) with the major contribution to the proportional increase arising from subclusters 0,1, 5, and 6 (**Fig.5E; Fig.S5D**). These macrophage subclusters were identified as tumor associated macrophages (TAMs) based on high expression of *Adgre1*, *Cd68*, *Csf1r*, *Itgam*, *Mertk*, and *Trem2* (**Fig.S5E**).

### The Omental Niche Drives a Pro-Inflammatory TAM Program

We further classified TAMs into SPMs (expression of *Ly6c2, Ccr2, Cd14, Fcgr1*) and LPMs (expression of *Adgre1, Timd4, Gata6, Icam2*) ^37,38^(**Fig.5F**). Of note, LPMs represented the dominant macrophage population in naïve peritoneal lavage (**Fig.5G**) and are subsequently more likely to be the first population encountered by DTCs in the peritoneal cavity. To better understand similarities and differences between LPMs and SPMs at different niches, we performed RNA velocity analysis. We observed a dynamic transition from ascites-derived SPMs to omental SPMs (cluster 1 to 0), and ascites-derived LPMs to omental LPMs (cluster 5 to 6) (**Fig.5H**). We then assessed the niche-specific functional states of resident LPM in ascites (Asc.LPMs) and metastatic omenta (Om.LPMs) (**Fig.5I**). Asc.LPMs were enriched in tissue residency markers (*Cmklr1*^39^) and components of MHC class I (*B2m, H2-T23*) while Om.LPMs activated programs related to NF-κB signaling (*Nfkbiz, Tnfaip3*), costimulation (*Cd86*), and chemokines (*Cxcl2*) (**Fig.5I**).

Given the apparent macrophage niche-dependent differentiation states, we next asked whether omental TAMs reflect a polarized, terminal state within the metastatic niche. If so, increased TAM polarization would likely shape the T cell compartment^40,41^. To assess this, we first subsetted macrophages (*Cd68*^high^/*Adgre1*^high^, **Fig.S5F**) and scored them against M1 and M2 gene modules. Compared to ascites-derived TAMs, omental TAMs exhibited higher hybrid functional state scores (**Fig.S5G**), reflecting an immunomodulatory TAM cellular state^42^ that cannot be captured by binning into a strictly immune-promoting or suppressing category. Unlike the immunomodulatory and ECM-related omental signatures, ascites-derived TAMs were enriched in pathways associated with cellular stress responses and NRF2-dependent redox metabolism (**Fig.5J & 5K**). On the gene level, ascites TAM signatures were reflected in elevated expression of key stress-related genes – including *Anxa1* and *Hmox1* – as well as components of mitochondrial respiration*, Ndufb8* and *Cox5a* (**Fig.S5H**). Indeed, while the ascites-derived T/NK compartment was dominated by a mixture of naïve CD4+ and CD8 +T cells, omental metastases and niche tissue contained a broader and more complex repertoire of activated, exhausted, and regulatory T cells (**Fig.5L**) enriched in pathways related to regulation of immune cell activation (**Fig.S5I**). Flow cytometry analysis at metastatic endpoint corroborated higher CD8 T cell infiltration in metastatic omenta compared to ascites (**Fig.S5J**). These findings indicate that ascites TAMs may represent an earlier, less polarized yet metabolically active state, while omental TAMs have transitioned into an immunomodulatory, polarized phenotype.

### Peritoneal Macrophages are Direct DTC Pro-survival Interactors

We first tested whether macrophages are functionally important in promoting ovarian cancer metastasis. Macrophage depletion via direct i.p. liposomal chlodronate (chlodrosome) injections (**Fig.6A**) demonstrated that peritoneal TAMs are pro-metastatic: depletion significantly reduced omental burden, as evidenced by reduced omental weight, tumor area across tissue sections, and reduced CD45- cancer cell frequencies in ascites at experimental endpoint (**Fig.6A-6D; Fig.S6A** gating strategy). Although chlodrosome treatment targets broad macrophage populations, we noted a preferred depletion of ascites LPMs, leading to a 51% reduction of this population at metastatic endpoint (**Fig.S6B**).

**Figure 6:**
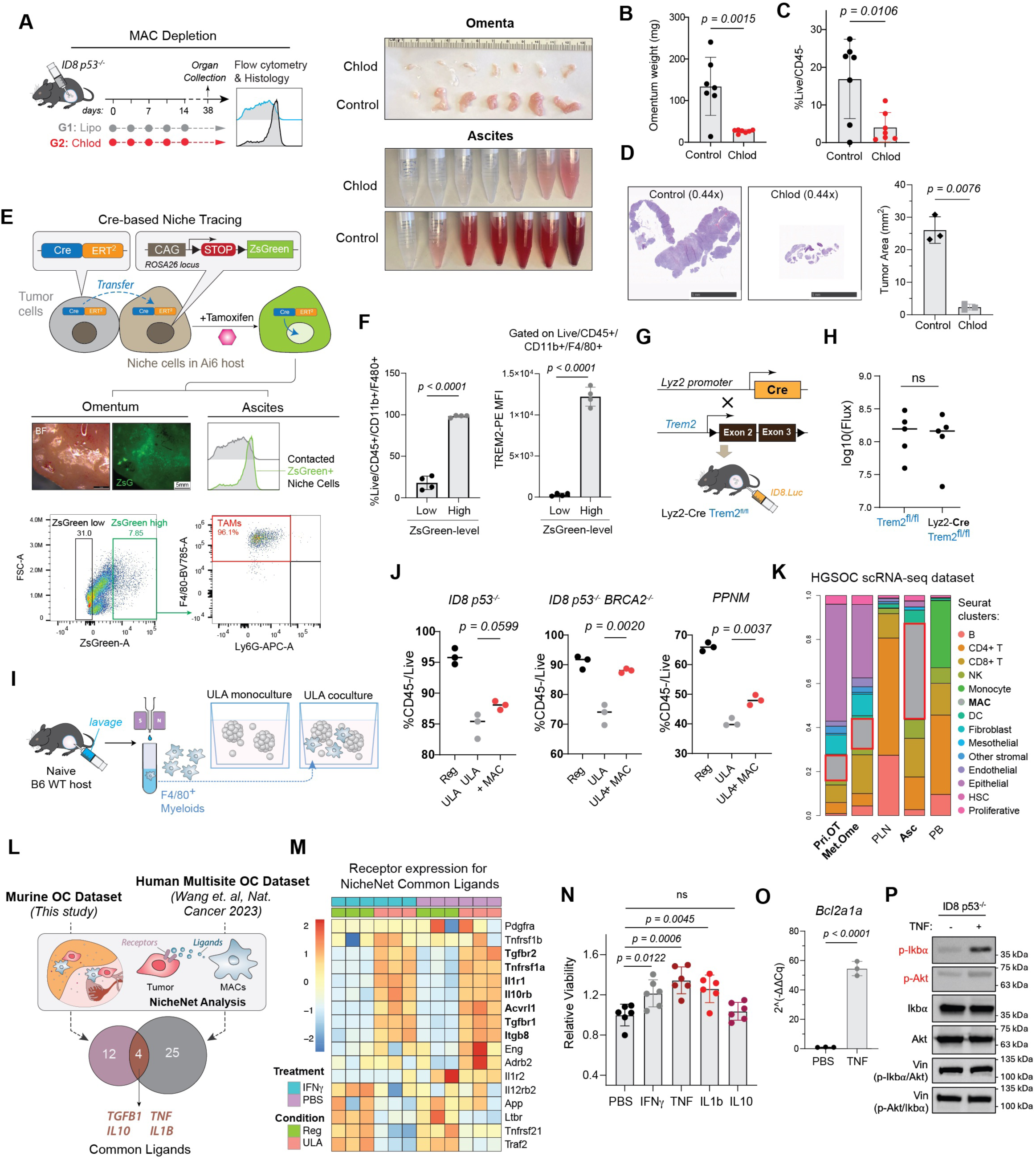
Peritoneal Macrophages Sustain DTC Fitness by Mediating TNF-induced Pro-survival Signaling. (**A**) Liposomal chlodronate-based macrophage depletion strategy and timeline schematic with representative tissue images. Omental tumor burden quantification (**B**) and flow-based ascites quantification (**C**) for each sample (n = 7 mice per group). (**D**) Representative H&E images taken at a fixed magnification (left) and tumor area quantification for control and chlodrosome-treated mice (right). For each group, three blocks were sectioned at 100 um intervals, and the average tumor area of 5 intervals was taken per datapoint. (**E**) Schematic of Cre-based NicheTracing system where ID8 and E0771 cancer cells express Cre.ERT2. Upon nuclear entry of Cre.ERT2 into Ai6 reporter cells, labeled cells turn ZsGreen positive (top). Representative gating strategy depicting LPMs as the major cell population tagged with zsG (bottom). (**F**) Quantification of F4/80 and TREM2 expression (MFI) in contacted, ZsG^high^ cells (n =4 tamoxifen treated Ai6 mice). Flow cytometry data was plotted as mean with SD and differences between groups were compared via a two-tailed unpaired Student’s t test. (**G**) Schematic depicting transgenic mouse model generation for macrophage-specific Trem2 knockout. (**H**) Log transformed BLI-derived flux values in Trem2fl/fl (wild type; n = 5) and Lys2-Cre/Trem2fl/fl (knockout; n = 5) mice at metastasis endpoint. Mice were injected with ID8 p53^-/-^ Luciferase cells. (**I**) Schematic of peritoneal macrophage isolation from naïve mice and macrophage-cancer cell coculture setup. (**J**) Plots of percent CD45 negative cancer cell viability (ZombieAqua negative) post coculture in three cell lines, n= 3 samples per group. Data was analyzed using a two-tailed Student’s t test and plotted with median values. (**K**) Stacked bar chart of main cell type proportions captured in human scRNA-seq dataset (Mendeley Data^69^) across five anatomical locations, using maintype-2 annotation provided by publisher. (**L**) Venn diagram of top NicheNet predicted ligands in ascites after filtering for robust expression in macrophages and overlapping with top ligands from murine dataset myeloids. (**M**) Bulk RNA-seq heatmap comparing the enrichment of NicheNet-related receptors in ULA-cultured cells treated with IFNγ compared to attached (Reg) culture conditions. RNA-seq data was analyzed using DESeq2 and visualized as variance-stabilized counts. (**N**) Cell-Titer Glo (CTG) assay luminescence assay graphed as fold change for ULA culture, with and without cytokine treatment, and comparing viability to the PBS-treated group. Fold change for each biological replicate (n = 6) was determined by normalizing the cell line’s CTG value to the regular attachment, without cytokine treatment, baseline. CTG assay luminescence data was analyzed using a two-tailed Student’s t test and graphed as mean with SD. (**O**) RT-qPCR derived Bcl2a1a transcript levels in ID8 p53^-/-^ cells under PBS and TNF treatment, n = 3 per group where each dot represents the average of technical replicates. Fold change (2^(-ΔΔCq)) data was analyzed using a two-tailed Student’s t test and graphed as mean with SD. (**P**) Western blot probing for p-Ikbα/Ikbα and p-Akt/Akt protein levels in ID8 p53^-/-^ cells under ULA culture, with or without TNF treatment.

To identify cells that have been in close contact with ovarian metastases, we developed a non-SP-TATK Cre-based iteration of the “NicheTracing” system^43^ to genetically trace microenvironmental contacts *in vivo* over a prolonged timeframe. In this system, cancer cells expressing Cre.ERT2 were injected into Ai6 (RCL-zsGreen) Cre reporter mice. Upon intercellular-cytoplasmic transfer and Tamoxifen-mediated nuclear entry of Cre into neighboring cells, Cre-mediated recombination turns on the zsGreen cassette in host cells (**Fig.6E**). After validating Cre.ERT2 expression in transduced cell lines (**Fig.S7A**), we benchmarked this system using *in vitro* coculture (**Fig.S7B-S7F**) and in an independent model of TNBC lung metastasis established using the E0771 cell line (**Fig.S7G-H**). In the ovarian metastasis model, we readily detected zsGreen^high^ (ZsG^high^) niche cells in both metastasis-bearing omenta and ascites at the experimental endpoint six weeks post cancer cell injection (**Fig.6E**). We performed flow cytometry analysis on ascites and gated cells into ZsG^low^ and ZsG^high^ (**Fig.6F**). Unlike lung metastases where only ∼30% ZsG^high^ cells were F4/80^high^ macrophages (**Fig.S7H**, right panel), over 90% of ZsG^high^ host cells were F4/80^high^ LPMs (**Fig. 6F**). Collectively, the above data suggest that LPMs interact with tumor cells in ascites and have pro-metastatic functions.

We next explored two mechanistic angles to determine how TAMs interact with DTCs. First, we observed that a significant portion of ZsG^high^ F4/80^high^/Ly6G^low^ TAMs expressed the efferocytotic marker Trem2 (**Fig.6F; Fig.S7I & S7J**). As Trem2 macrophages are known to play lipid sensing, anti-inflammatory and immunosuppressive tumor promoting roles^44–46^, we specifically analyzed Trem2^high^ and Trem2^low^ TAM populations in our scRNA-seq dataset (**Fig.S7K**). Trem2^high^ cells showed high expression of broad chemotaxis, ERK pathway, and adhesion pathways (**Fig.S7L-S7N)**. In contrast, significantly enriched pathways in Trem2^low^ cells were related to epithelial cell proliferation, mesenchymal transition, and TGFβ pathways (**Fig.S7N**), indicating distinct roles. After induction of experimental ovarian metastasis in macrophage-specific (Lyz2-driven) Trem2 KO mice (**Fig.6G; Fig.S7O & S7P**), unexpectedly, we observed no significant effect of Trem2 KO on ovarian metastasis burden (**Fig.6H**). As ablating Trem2 signaling in macrophages was not sufficient to change the overall ovarian cancer metastasis progression, this suggests that peritoneal macrophages could promote metastasis independently of known Trem2-mediated efferocytotic and anti-inflammatory programs.

Given that the predominant niche contacts *in vivo* are established with LPMs, the enriched cell type after early DTC priming, and having ruled out the Trem2 efferocytotic program, we hypothesized that peritoneal macrophages instead directly interact with cancer cells to promote initial colonization events. Although previous studies have reported that M2-polarized TAMs could promote tumor cell spheroid formation^47^, it remains unclear if and how naïve peritoneal macrophages could promote early DTC fitness. We performed a macrophage-cancer cell coculture under ultra-low attachment (ULA) conditions (**Fig.6I**). Naïve macrophages from peritoneal lavage were enriched using F4/80 magnetic beads (**Fig.S6C**, flow gating strategy) and cocultured with three different murine ovarian cancer cell lines; ID8 p53^-/-^, ID8 p53^-/-^ BRCA2^-/-^, and PPNM^48^ (**Fig. 6J; Fig.S6D** flow gating strategy). After a 48-hour coculture period, flow cytometry analysis revealed that cancer cell viability, as measured by live/dead staining (ZombieAqua), decreased under ULA conditions compared to regular attachment plates. Most importantly, survival was partially rescued with macrophage coculture across all three cell lines (**Fig.6J**). These results indicate that peritoneal macrophages directly support cancer cell survival under low attachment conditions.

To determine whether this pro-survival macrophage-DTC interaction is conserved in human ovarian cancer, we analyzed a human-derived single-cell dataset containing over 200,000 single cells from 14 ovarian cancer patients collected from five locations: peripheral blood (PB), peripheral lymph nodes (PLN), malignant ascites (Asc), primary tumor (Pri.OT), and matched metastatic tumor (Met.Ome)^49^. As observed in our murine scRNA-seq dataset, cancer cell-containing compartments, including Asc/Pri.OT/ Met.Ome, had increased proportions of macrophages compared to PB and PLN (**Fig.6K; Fig.S8A**). DEG analysis between TAMs at metastatic sites (**Fig.S8B**) and subsequent GO pathway analysis conducted on the DEG list revealed immune-modulatory pathways (cytokine, chemokine, allograft rejection) were upregulated in Om.TAMs while cargo receptor activity related genes were enriched in Asc.TAMs (**Fig.S8C**).

### Macrophage-derived cytokines rescue ascites DTCs from anoikis

We applied NicheNetR analysis to further characterize ligand-receptor interactions between macrophages (ligand provider) and tumor cells (signal receiver through receptor) (**Fig.6K**). We implemented this strategy in both our murine dataset (**Fig.S8D**) as well as the patient dataset (**Fig.S8E**). After defining myeloid cells as the sender cell type in the murine dataset and macrophage cluster as the sender in the human dataset, we compared the top predicted tumor-interacting ligands in each dataset. We identified four overlapping sender ligands across species - *TGFB1*, *IL10*, *TNF, and IL-1β* - that were the most significant ligands predicted to interact with cancer cell receptors (**Fig.6L, Fig.S8D & S8E**). These ligands were expressed at high levels in patient macrophages (**Fig.S8F**). Next, we examined the top predicted downstream target genes in cancer cells affected by top TAM ligands. Pathway analysis of the predicted downstream genes yielded pathways involved in cell-cell adhesion, response to oxygen levels, and apoptosis regulation (**Fig.S8G**). Importantly, both SPMs and LPMs in our *in vivo* murine scRNA-seq dataset expressed the four common ligands (*TGFB1, IL10, TNF, and IL-1β*). We next derived a list of possible corresponding receptors to these four ligands based on “prior interaction potential” in the NicheNet analysis. Interestingly, cross-examination of corresponding common receptor expression in the bulk RNA-seq data derived from ULA culture condition (**Fig.4** RNA-seq panels) revealed a subset of up-regulated receptors specific to the ULA culture condition (e.g. *Tgfbr1/2*, *Il1r1*, and *Il10rb*). Such ULA-dependent receptor up-regulation was largely independent of IFNγ treatment (**Fig.6M**). In parallel to cancer cell intrinsic IFNγ programs, these data suggest that TAMs provide a cocktail of ligands that could potentially amplify pro-survival signals in IFNγ-primed DTCs.

We next assessed the potential of TNF, IL10, and IL-1β cytokines to promote anchorage independent survival *in vitro*. TNF and IL-1β, but not IL10 treatment, significantly increased CTG flux in an anchorage-dependent manner (**Fig.6N; Fig.S8I**). TNF has been reported to promote survival pathways in contexts where the TNF receptor remains sequestered on the cell surface upon TNF binding, and surface sequestration can occur through receptor sialylation via the St6gal1 sialyltransferase^50^. In line with that notion, we observed increased expression of the anti-apoptotic gene *Bcl2a1a* after TNF treatment (**Fig.6O**), and ULA culture primed cancer cells to express higher levels of *St6gal1 in vitro* (**Fig.S8J**). Mechanistically, TNF treatment induced the activation of canonical pro-survival pathways, including NF-κB (measured via p-Iκbα) in three cell lines and Akt (measured via p-Akt) in two cell lines (**Fig.6P; Fig.S8K**).

### A Combined DTC Pro-survival Signature Predicts Clinical Outcome in Ovarian Cancer

In this study, we identified two convergent pro-survival programs that enable DTC fitness in the peritoneal cavity: a tumor-intrinsic IFNγ response and a macrophage-derived paracrine signaling collectively protect DTCs from anoikis. To assess whether these pathways carry clinical relevance, we constructed a composite gene signature derived from our findings (25 genes related to DTC IFNγ signature & 4 genes related to macrophage-DTC NicheNet analysis) for prognostic potential using multivariate Cox proportional hazard modeling across multiple ovarian cancer patient cohorts (**Fig.S9A**).

For overall survival (OS), *Ifngr1* was the largest contributor to the risk score, consistent with our data demonstrating that loss of IFNγ receptor signaling significantly reduced metastatic burden and extended survival in the mouse model (**Fig.S9A**). In addition, *St6gal1*, a TNF-responsive gene implicated its pro-survival signaling, ranked as the second highest contributor to OS risk (**Fig.S9A**). For recurrence-free survival (RFS), IFNγ-inducible *Gbp2b/GBP1* has the highest risk score alongside with *Marco* and *Stat1* (**Fig.S9A**). We next stratified patients using our 29 gene composite gene signature median risk score. In both discovery cohorts and independent validation cohorts, ovarian patient tumor gene expression with a high-risk score showed a consistent reduced OS and RFS (**Fig.S9B & S9C**). These results indicate that the collective contribution of DTC-intrinsic and macrophage-derived pro-survival programs characterized in this study jointly carry prognostic significance in HGSOC.

## DISCUSSION

### Early DTC Dominance Phenotype in Ovarian Cancer Metastasis

DTC-niche interactions at each stage of the metastatic cascade fundamentally determine disease progression and patient outcomes. Following the initial wave of metastatic dissemination, early DTCs inevitably reshape the metastatic niche microenvironment, irrespective of whether they ultimately survive or perish upon arrival. While the pre-metastatic niche (PMN) has been extensively characterized^51,52^, the dynamic processes following DTC seeding represent a less explored system, which we term the Post-Seeding Metastatic Niche (PSMN). The multi-cellular dynamics resulting from progressive DTC accumulation at newly colonized PSMN mirrors a well-established ecological concept: the Allee effect, which describes how population growth rates depend on population density through competition or cooperation. Although the Allee effect has been demonstrated in metastatic contexts using *in vitro* models^53^, how continuously arriving DTCs dynamically alter PSMN properties and influence the fate of later-arriving tumor cells within the intact physiology of living systems remains unstudied.

Leveraging our MetTag barcoding and staggered injection model, the data presented here consistently demonstrated a skewed overrepresentation of early DTCs: TP-1 and TP-2 DTCs overwhelmingly dominate the metastatic burden at day 40, whereas cells introduced at later days (TP-6) showed relatively reduced contributions to final overt metastases. This pattern persists across immunocompetent, immunodeficient (NRG), and macrophage-depleted hosts (**Fig.1**). This skewness in early DTC dominance cannot simply be attributed to inherent, longer chronological growth times of early DTCs, as there remains an ample growth window for TP-6 cells (from Day 11 to Day 40). Additionally, these results are consistent with early cancer cell dissemination suggested by clinical tumor genetics studies in ovarian and breast cancer metastasis^11–17^. There are several non-mutually exclusive hypotheses that could explain early DTC dominance observed in our study. First, the omentum might contain a limited number of permissive micro-niches that favor DTC colonization. Pioneer DTCs might rapidly saturate these micro-niches and monopolize spatial resources – a relative “first come, first served” model operating on a shorter time scale of tumor evolution. Second, the immune-independent nature of skewed distribution of temporally disseminated DTCs suggests a role of non-immune stromal elements in dictating colonization success. Early DTCs may actively remodel the stromal niche, rendering it less favorable for later DTCs. In addition to these inhibitory mechanisms, early DTCs may in parallel build a niche to support themselves and, to a lesser extent, later arrivers as their complete absence abolishes later metastatic outgrowth (**Fig.3L-M**). Comprehensive spatiotemporally resolved imaging and time course analyses are required to further explore these hypotheses.

### Duality of IFNγ Signaling-driven Metastatic Adaptation

Beyond the “seed and soil” hypothesis, it has been increasingly appreciated that metastasis is not only not merely passive fitness-based selection of DTC clones at the niche, but also an active epigenetic DTC-niche mutual adaptation process. Our single-cell transcriptome analysis, coupled with barcoding clonal analysis, provided us with clonal resolution of the underlying genetic and epigenetic determinants governing DTC behavior. Interestingly, instead of occupying unique transcriptome clusters - which would likely suggest distinctive genetic mutational features - the expanded clones form similar coordinated adaptive “mini-ecosystems” encompassing multifaceted transcriptome features that converge toward an inflammatory/IFNγ response (**Fig.2H-L**). This data aligns with single-cell whole-genome sequencing (scWGS) of treatment naïve ovarian cancer patients which details the existence of (1) significant clonal heterogeneity within the same patient and (2) distinct IFNα/IFNγ expressing clones^54^.

Our CRISPR/Cas9 *in vivo* screen results (**Fig.3**) further support the notion that cancer cell fitness in the peritoneal cavity and subsequent omental metastasis depend on rapid, transcriptional-level adaptation to IFNγ-responsive programs. As previously noted, our data indicate that the early DTCs IFNγ response is required for subsequent metastatic seeding (**Fig.3L-N**). Knocking out Ifngr1 in the early batch of DTCs significantly reduced the metastatic potential of later DTCs with intact Ifngr1 (as evidenced by total metastatic burden), indicating a critical temporal window in which early IFNγ responsiveness and active PSMN conditioning by initial colonized DTCs jointly drive the subsequent metastatic continuum. One important hypothesis to consider in relation to these results is the influence of classical concomitant tumor immunity. This framework describes a phenomenon where a primary tumor restricts distant metastases in an antigenically matched, T cell-dependent, manner^55,56^. While the failure of Ifngr1-KO pioneer DTCs to efficiently colonize may result in an increase of available antigens for host immune cell priming supporting this framework, the persistence of early DTCs in immunodeficient NRG mice (Fig.1) suggests a more fundamental principle for the observed seeding bottleneck. Our findings expand the conventional view of clonal fitness by revealing its temporal and niche-dependent nature, where transient adaptive states at critical windows determine metastatic success. In this “first come, first served” model, early metastatic DTC–niche crosstalk through IFNγ sensing-mediated metabolic and epigenetic adaptation determines metastasis success by enabling the earliest colonizers to occupy and remodel permissive micro-niches while restraining later DTCs’ entry.

Although the findings presented here support a tumor cell-centric pro-metastatic role of IFNγ, IFNγ in metastatic niches can act in complex ways. In some contexts, it reduces metastasis through activation of effector T and NK cells, thereby enhancing the effectiveness of immunotherapy^57,58^, while in other contexts IFNγ can induce apoptosis in regulatory T cells and inhibit angiogenesis^59^. Specifically in ovarian cancer, both anti-metastatic and pro-metastatic roles have been reported^60,61^. IFNγ has been shown to synergize with beta glucan treatment to reduce peritoneal metastasis and Ifngr1 knockout in host cells leading to a significant increase in metastasis^62^. For pro-metastatic roles, IFNγ has been previously reported to upregulate PD-L1 and IL-8 in ovarian cancer cells, leading to increased cell migration and proliferation^61^. We, however, did not observe any proliferation differences via colony formation assay *in vitro* (**Fig. S4A**) after a 7-day IFNγ exposure. Given that *Cd274* and *Psmb10* transcript levels were not increased in Irf1^high^ cells *in vivo* (**Fig.4J**), we focused on *Parp14* induction as an active pro-survival cellular adaptation maintained in ascites over time, as opposed to an immediate physiological response to IFNγ signaling. Based on the data presented here, we believe that, although both effector immune cells and cancer cells could be impacted by IFNγ, IFNγ response in cancer cells (intrinsically) leads to an overall pro-metastatic effect. This effect is an important mechanism driving metastasis and it should be carefully considered when designing future anti-metastasis treatment. Importantly, we do not propose that Ifngr1 downstream signaling in cancer cells itself directly reshapes the omental niche. However, we do observe that the loss of attachment conferred by the ascites microenvironment: (1) affects downstream interferon signaling which impacts survival and (2) impacts surface expression of receptors responsible for tumor-macrophage interactions.

### Macrophage Plasticity and Pro-survival Ligands

Cells of the myeloid lineage are characterized by significant plasticity, especially in disease^63,64^. However, recent studies point to niche-regulated plasticity restriction^65^. Upon infiltration of inflammatory monocytes into metastatic sites, they often exhibit “commonly described” reprogrammed states such as increased checkpoint expression or efferocytotic receptor upregulation due to cancer cell engulfment. In addition to these classical reprogramming events, myeloid cells exhibit tissue-specific reprogramming after prolonged residence at a specific niche^65^. Thus, although we observe Trem2^high^ SPMs and LPMs in metastatic omenta and ascites, depleting Trem2 signaling did not significantly impact the metastasis phenotype (**Fig.6H**), pointing the role of other programs within the broad LPM population that could provide support to cancer cells. Although IFNγ treatment is sufficient to trigger expression of anti-apoptotic genes such as Parp14, and loss of attachment triggers IL-1β, TNF, TGFβ, and IL10 receptor transcription (**Fig.6M**), our data suggests that macrophages (extrinsically) provide multiple pro-survival signals to these primed cancer cells. Specifically, TNF and IL-1β are known amplifiers of NF-κB^66,67^, a master-regulator of anti-apoptotic and anti-necrotic gene expression^68^. Given our *in vitro* findings with TNF and IL-1β treatment (**Fig.6N**), we hypothesize that initial IFNγ exposure and loss of attachment prime DTCs to respond strongly to macrophage-derived pro-survival cues and collectively amplify pro-survival programs in DTCs. Importantly these findings build upon prior knowledge on the role of a EGF/EGFR paracrine signaling loop between tumor cell and TAMs in promoting adhesion of cancer cells through integrins, and thereby spheroid formation^47^.

In summary, our study dissected the complex landscape of ovarian cancer metastasis through a systematic characterization of both cancer cells and local immune factors using MetTag temporal, clonal, and scRNA-seq analysis. Our findings highlight the importance of ascites as a functional reservoir for pro-survival factors and that early acquisition of IFNγ response and interaction with macrophages are necessary deterministic adaptation steps for DTCs to survive and colonize.

## STUDY LIMITATIONS

While our *in vivo* studies using murine ID8 p53^-/-^ cells allowed us to explore cancer cell-immune interactions in a fully immunocompetent background, we acknowledge its differences from human HGSOC histologic features, disease heterogeneity, and complexity of genomic alterations. Further studies in this area would benefit from including other experimental models of ovarian cancer. Additionally, at advanced stages of ovarian cancer modeled in this study, a significant contribution to the ascites cancer cell fraction likely arises from secondary seeding from established omental implants which we cannot distinguish using our barcoding strategy. Furthermore, the CRISPR screen gene list was restricted to only the top DEGs due to in vivo coverage constraints and served to dissect the main pathway modules, as opposed to identifying the most significant single gene driver. Dropout-based CRISPR screening approaches are inherently biased towards genes that are necessary for both cell survival and later proliferative expansion. IFNγ responsive genes may facilitate survival in ascites without creating a sufficiently large magnitude of cell proliferation differences to yield strong dropout hits. Consistent with our interpretation, several IFNγ responsive genes (Marco and Gbp2b) had higher false discovery rate (FDR) values in ascites, which we interpreted as a technical limitation of dropout screening in slower cycling cells. Importantly, although we focused on the role of IFNγ signaling in anoikis resistance, pro-metastatic effects of IFNγ likely extend beyond these survival pathways. Finally, while we utilized NicheTracing to assess cancer cell and immune cell interactions in ascites, our study lacks (1) comparative tracing results in the omental niche and (2) real-time imaging or tissue level immunofluorescent staining of DTC interactions at different time points.

## ACKNOWLEDGMENTS

This study was supported in part by NIH training grant T32GM075762 (to Aleksandrovic); NIH grants R01 CA194697-01, R01 CA255064-01A1, and R21 CA263798-01 (to Zhang); NIH grant R35GM142654 (to Zhong); NIH grants RM1GM145399 and U54CA268072 (to Dean); Cancer Prevention and Research Institute of Texas (CPRIT) Scholar Award RR220024 (to Zhang); CPRIT Individual Investigator Research Award RP230261 (to Zhong); and CPRIT Core Facility Grant RP250571 (to Dean). We additionally acknowledge support from the Dee Family endowment (to Zhang). We thank the following core facilities for their support: the Notre Dame Freimann Life Sciences Center, the Notre Dame Genomics and Bioinformatics Core Facility, the Indiana University School of Medicine Center for Medical Genomics, the Indiana University Simon Cancer Center, the Simmons Comprehensive Cancer Center, the UT Southwestern Animal Resource Center, the UT Southwestern Flow Cytometry Core, the UT Southwestern Whole Brain Microscopy Facility, the UT Southwestern Texas Cancer Cell Imaging Core, and the UT Southwestern Small Animal Imaging Core Facility. The computational component of this project was supported in part by the BioHPC high-performance computing facility at UT Southwestern.

## AUTHOR CONTRIBUTIONS

Conceptualization and experiment design, E.A., M.S.S., J. Lea, and S.Z.; Data analysis, E.A., S.Z., X.L., Z. Zhao, H.M.B., A.T., K.M.D., L.Y., and L.X.; Experiments and methodology, E.A., S.Z., S.R.F., S.M.G., J. Lopez, W.M., N.D., T.C.R., M.Z., Z. Zhong, H.M.,B., K.M.D.; Manuscript writing and revision, E.A., S.M.G., and S.Z.; Study supervision, S.Z. We attest that all coauthors reviewed and pre-approved the manuscript before submission.

## SUPPLEMENTAL INFORMATION (Including Supplementary Figures S1-S9)

**Figure S1. MetTag Tracing Construct and Metastatic Model Validation**

(**A**) Schematic depicting MetTag library barcoding plasmid, and modes of detection upon plasmid integration: DNA level, Pol III U6 driven RNA level, and Pol II CROP-seq based. (**B-C**) Barcode frequencies comparing relative evenness of library distribution across BC.ID batches (B) and original depositor LARRY library (C). (**D**) Table depicting unique reads detected in each library batch after MiSeq sequencing and read filtering. (**E**) Representative colony formation assay wells for each MetTag-LARRY-BC.ID ID8 p53^-/-^ line. (**F**) Quantification of per well colony area (n = 3 per sample). Colony areas were compared in GraphPad Prism and plotted as mean with SD and differences between groups were compared via a two-tailed unpaired Student’s t test. (**G**) Representative ascites images and dissecting scope images of GFP+ metastases formed under the repeated injection schema in wild type (n = 4) and NRG mice (n = 4). (**H**) Separate cohort of mice (n = 5 per group) were subjected to ID8 p53^-/-^ Luciferase injection and omental burden was quantified via ex vivo BLI imaging (left) and by measuring omental weights (right). (**I**) Representative ascites images and dissecting scope images of GFP+ metastases formed under the repeated injection schema in liposome (n = 4) and chlodrosome treated mice (n = 3). (**J**) Bar chart depicting macrophage depletion efficiency (left) and quantification of GFP+ cells (right) in ascites under macrophage depletion. (**K**) RT-qPCR based assessment of DTR transcript expression in each MetTag-LARRY-BC.ID ID8 p53^-/-^ line after transfection with DTR or stuffer constructs. Flow cytometry and qPCR data was analyzed in GraphPad Prism and plotted as mean with SD and differences between groups were compared via a two-tailed unpaired Student’s t test. (**L**) Representative colony formation assay wells for each MetTag-LARRY-BC.ID ID8 p53^-/-^ line without (left) or with DT treatment (right).

**Figure S2. scRNA-seq-based Tumor Cell Identification and Cancer Cell Velocity**

(**A**) Feature plot of main marker genes used to define and subset cancer cells. (**B**) Split UMAP depicting cluster distributions across the four experimental conditions and identifying cancer cell-enriched clusters in ascites and omentum. (**C**) Reclustered cancer cell UMAP colored by the two metastasis-bearing anatomical locations. (**D**) Heatmap of reclustered cancer cell marker genes. (**E**) Violin plots of escape R generated Hallmark pathways each represented by a different color. Details on escape R ssGSEA can be found in the Methods section. (**F**) Log10-transformed frequency distributions of top 100 LARRY barcodes with counts >= 10, obtained from *in vitro* cell culture. (**G**) UMAPs depicting main Hallmark pathway signatures represented in the top three clonally expanded barcoded cells. (**H**) CellRank-generated initial and terminal cellular states overlayed on the cancer cell UMAP. (**I**) Violin plot of IFNα signature score across the six different BC.ID timepoints. IFNα signature was assessed by first defining the signature using Hallmark IFNα response genes, followed by directly applying the wilcox.test() function in Seurat to compare expression level differences between TP-1 and TP-6 groups. (**J**) UMAP of captured single cells in TNBC lung metastasis model (left) and RNA-based gating strategy to subset non-immune cells for subsequent barcode analysis. (**K**) Heatmap depicting enriched clonal LARRY barcodes in each lung sample (HTO n’s = 3).

**Figure S3. IFN Signaling in Metastatic Niches and *In vivo* CRISPR Screening**

(**A**) UMAP of captured cancer cells colored by *in vivo* (Lung.Met; HTO n’s = 2) and *in vitro* (E0771.cell; HTO n’s = 1) cell line. (**B**) ORA analysis-derived Hallmark pathways enriched *in vivo* compared to *in vitro*. Module scores comparing canonical IFNγ (**C**) and IFNα (**D**) genes between the two conditions. (**E**) VlnPlots of select individual IFNγ response related genes. (**F-G**) ORA analysis-derived pathways based on DEGs between AscMet and OmMet. (**H**) Heatmaps depicting individual IFN response gene scoring differences between AscMet and OmMet. (**I**) Western blot validation of SpCas9 expression in ID8 p53^-/-^ cells after PiggyBac transposon transfection and selection with hygromycin. (**J**) Representative flow cytometry generated histogram of mCherry fluorescence (left) and bar chart depicting percent mCherry positive cells five days after lenti-guide transduction (right). (**K**) scRNA-seq derived violin plot of Ifngr1 expression levels in cancer cells *in vivo*. Module scores and individual gene transcript levels were compared using the wilcox.test() function in Seurat. (**L**) Representative flow cytometry histogram of IFNGR1 surface expression in knockout vs. wild type cells (left) and bar chart with quantification of IFNGR1 mean fluorescent signal changes (right). n = 3 biological replicates per group were analyzed using a two-tailed Student’s t-test and graphed as mean with SD. (**M**) Western blot probing for IFNγ downstream transcriptional targets, Stat1 and Gbp2b, in Ifingr1 (sg#1) knockout cells. (**N**) Endpoint BLI flux images of sgIfngr1 (n = 7) compared to sgNTC experiment mice (n = 8).

**Figure S4. IFNγ Downstream Mediators of Anoikis Resistance**

(**A**) Colony area-based quantification of clonogenic assays in absence and presence of recombinant IFNγ, with or without Ifngr1 (sg#1) knockout. Colony area was plotted in GraphPad Prism as mean with SD and analyzed using a two-tailed Student’s t test. (**B**) Schematic depicting *in vitro* bulk RNA-seq experiment comparing ULA and regular culture conditions under IFNγ treatment. (**C**) PCA plot of sequenced RNA-seq groups, generated from variance-stabilized transformed (VST) counts. Each dot represents a different sample; color represents different treatment or culture condition. (**D**) MA quality control plot showing DEGs between ULA and regular culture. Blue points indicate significant genes (padj < 0.05, log2FC>+/- 1). (**E**) S and G2/M cell cycle gene heatmap comparing wild type sgNTC cells cultured under ULA and regular culture conditions. (**F**) Heatmap of top DEGs - both unique and overlapped with regular culture - between PBS and IFNγ treated cells under ULA culture (n = 3 per condition). (**G**) Western blots showing partial knockout of Stat1 and Irf1 transcription factors, as well as subsequent ablation of Gbp2b expression with IFNγ treatment, as a proxy for downstream signaling ablation. (**H**) Violin plots comparing expression levels of immunoproteasome (Psmb10, Psmb9), DNA repair/apoptosis (Parp12, Parp14, Parp9), and immune checkpoint (Cd274) levels between OmMet (n = 2) and AscMet (n = 4). (**I**) Violin plots comparing expression level of Cd274 (left) and Psmb10 (right) between Irf1^high^ and Irf1^low^ cells. scRNA-seq based transcript expression levels depicted in (H) and (I) were compared by applying the wilcox.test() function in Seurat. (**J**) Western blot depicting partial knockout of Parp14 in ID8 p53^-/-^ cells.

**Figure S5. scRNA-seq Analysis of Myeloid and Lymphoid Populations**

(**A**) UMAP projection of all cells captured across four sequenced groups, consisting of two integrated datasets. (**B**) “Broad_cell.ID” classification of Seurat clusters into cell types based on cluster marker genes. (**C**) CellChat generated bar charts depicting number of interactions (left) and interaction strength (right) between different cell types across the four sample groups. (**D**) Heatmap of myeloid TAM subclusters 1, 0, 5, and 6. (**E**) Violin plots showing gene expression of canonical TAM markers across myeloid subclusters. (**F**) RNA-based gating strategy to subset out macrophages for subsequent analysis. (**G**) Biaxial scatter plot of M1 and M2 module scores with overlapped ascites and omental TAMs. (**H**) Violin plots comparing individual genes belonging to cellular response to stress (*Anxa1, Hmox1*) and cellular respiration (*Ndufb8, Cox5a*) pathways between omental and ascites TAMs. Hybrid score depicted in (I) and violin plot expression levels in (H) were compared using the wilcox.test() function in Seurat. (**I**) Top ORA-derived GO Biological Process pathways upregulated in omentum-derived compared to ascites-derived T/NK cells. (**J**) Flow cytometry analysis of CD8+ T cells frequencies in naïve and metastatic settings. Flow cytometry data was analyzed via two-tailed unpaired Student’s t test and plotted as individual values with median.

**Figure S6. Flow Cytometry Gating Strategies**

(**A**) Flow gating strategy for macrophage depletion study and representative gates for F4/80 macrophages in liposome control and chlodrosome treated groups. (B) Quantification of F4/80^high^ LPMs in control and chlodrosome treated mice. P value was generated using a two-tailed Welch’s t-test and corresponded to a 51% decrease in macrophage frequency. (**C**) Flow gating strategy for assessing MHC-II+/CD11b+ macrophage enrichment following F4/80 magnetic bead isolation from peritoneal cavity. Bar plot showing relative enrichment of macrophage populations before the start of coculture. (**D**) Flow gating strategy assessing CD45- cancer cell viability (Zombie Aqua negative population) after co-culture completion.

**Figure S7. Mechanistic Exploration of TAM Pro-metastatic Function**

(**A**) Western blot validation of Cre expression in target cell lines: E0771 murine TNBC line and ID8 p53^-/-^ ovarian cancer cell line. (**B**) Schematic depicting Cre reporter used to transfect HEK293T cells and Cre constructs expressed by target cells for *in vitro* color switch assays. (**C-F**) Fluorescence cell culture images generated after 72 hours of cancer cell-CreERT.2/HEK293T-reporter coculture with or without 2 uM 4-OHT treatment. (**G**) Representative dissecting microscope images of negative control spleen and lugs collected 2 weeks post E0771-CreERT.2 retro-orbital injection. (**H**) Flow cytometry bar plots at experimental endpoint showing percent switched immune cells (ZsG+) with (n = 4) or without (n = 4) tamoxifen treatment (left) and within the tamoxifen treated group, frequency of ZsG+ macrophages compared to ZsG- (right) in metastatic lungs. (**I**) Bar plot showing frequency of ZsG+ CD11b myeloids compared to ZsG- in malignant ascites (n = 4). Flow cytometry data in (H) and (I) was analyzed using a two-tailed unpaired Student’s t test and plotted as mean with SD. (**J**) Representative flow cytometry gating for TREM2 macrophage enrichment in ZsG+ fraction. (**K**) scRNA-seq gating strategy used to subset TREM2-high and low TAMs. (**L**) Volcano plot comparing DEGs between metastasis-infiltrating Trem2-low and Trem2-high macrophages. Significant DEGs were generated using the FindMarkers() function in Seurat and Wilcoxon test to compare gated Trem2-low and Trem2-high cells. (**M-N**) Go Biological Processes pathways upregulated in Trem2-high (left) and Trem2-low (right) TAMs. Macrophage-specific Trem2 depletion validation representative flow cytometry plots (**O**) and quantification (**P**).

**Figure S8. Patient and Murine Tumor-TAM NicheNet Analysis and Mechanism**

(**A**) UMAP depicting all (>200,000) single cells collected from five anatomical locations and annotated by cell type (“Maintypes-2”). (**B**) Heatmap of DEGs between Ascites and Met.Ome-derived macrophages. DEG analysis was performed using the FindMarkers() function in Seurat. (C) Dot plots depicting top GO molecular function pathways expressed in patient omental TAMs (left) and ascites TAMs (right). (D) NicheNet heatmap depicting top receptor-ligands pairs in murine ascites. (E) NicheNet heatmap depicting top receptor-ligands pairs in patient ascites. (F) Bubble plot of NicheNet-derived ligand expression among Maintypes-2 clusters. (**G**) Dot plot of GO pathways enriched in cancer cells based on regulatory potential and downstream genes affected by macrophage ligands. (**H**) Murine scRNA-seq violin plots showing transcript levels of *Tnf*, *Il1b*, *Il10*, and *Tgfb1* in SPMs and LPMs. Expression levels were compared using the wilcox.test() function in Seurat. (**I**) Cell-Titer Glo (CTG) assay luminescence assay graphed as fold change for Reg culture, with and without cytokine treatment, and comparing viability to the PBS-treated group. Fold change for each biological replicate (n = 6) was determined by normalizing the cell line’s CTG value to the regular attachment, without cytokine treatment, baseline. CTG assay luminescence data was analyzed using a two-tailed Student’s t test and graphed as mean with SD. (**J**) RT-qPCR derived St6gal1 transcript levels in ID8 p53^-/-^ cells under Reg and ULA culture conditions, n = 6 per group where each dot represents the average of technical replicates. Fold change (2^(-ΔΔCq)) data was analyzed using a two-tailed Student’s t test and graphed as mean with SD. (**K**) Western blot probing for p-Ikbα/Ikbα and p-Akt/Akt protein levels in ID8 p53^-/-^ BRCA2^-/-^ cells and PPNM cells under ULA culture, with or without TNF treatment.

**Figure S9. Prognostic Value of Interferon and NicheNet-based Gene Signatures in HGSOC Patients**

(**A**) Bioinformatics workflow using clinical RNA-seq datasets derived from the curatedOvarianData package and 29 genes from the present study (25 IFNγ-related and four NicheNet-derived genes). Genes were prioritized via multivariate Cox proportional hazard regression and iteratively selected from the datasets. (**B**) Kaplan-Meier plots depicting overall survival (OS) for patients stratified into high and low risk groups, based on the median risk score. Results are illustrated for a discovery cohort (left) and validation cohort (right). (**C**) Kaplan-Meier analysis of recurrence-free survival for patients based on risk score stratification in a discovery and independent validation cohort as described in (B). High and low risk groups were compared using a standard log-rank test.

## METHODS

### Experimental Models

#### Mice

All animal experiments were conducted according to IACUC protocols which were pre-approved by the University of Notre Dame and UT Southwestern Medical Center IACUC committees. Mice were housed with unlimited access to food and water in a temperature-controlled holding room with a standard 12-hour light/dark cycle. Health checks were performed daily during *in vivo* experiments. Wild type C57BL/6J (B6) mice were 8-10 weeks old at the experiment onset. Control and treated transgenic mice were age-matched, greater than 8 weeks of age. All experiments were carried out exclusively on female mice. C57BL/6J (B6, 000664), NOD.Cg-*Rag1^tm1Mom^Il2rg^tm1Wjl^*/SzJ (NRG, 007799), B6.Cg-*Gt(ROSA)^26Sortm^*^6^*^(CAG-ZsGreen^*^1^*^)Hze^*/J (Ai6, 007906), and B6J.129(Cg)-*Gt(ROSA)26Sor^tm1.1(CAG-cas9*,-EGFP)Fezh^*/J (Cas9 026179) mice were obtained from Jackson Laboratory (Ben Harbor, ME). We generated Lyz2-Cre(+/-)/Trem2(fl/fl) double transgenic mice by breeding Lyz2-Cre heterozygous (+/-) mice with Trem2(fl/fl). These lines were generous gifts from the Zhong Lab at UT Southwestern Medical Center.

#### Cell Lines

The TNBC E0771 cell line (CH3 Biosystems, Amherst, NY; Cat.No. 940001-Vial) was maintained in RPMI-1640 media supplemented with 10%FBS and 1% Penicilin/Streptomycin (Pen/Strep). ID8 p53^-/-^, ID8 p53^-/-^ BRCA2^-/-^, and PPNM ovarian cancer cell lines, provided by the Stack lab at the University of Notre Dame, were maintained in DMEM low glucose media with 5% FBS, 1% Pen/Strep, and 1% Insulin-Transferrin-Selenium (ITS). PPNM cell media was additionally supplemented with 100 ng/ml Cholera toxin and 2ng/mL epidermal growth factor (EGF). HEK293T cells were cultured in DMEM high glucose supplemented with 10% FBS, 1% Pen/Strep, and 1% GlutaMAX (Gibco, 35050061). Cell cultures were grown at 37 °C, 5% CO_2_, and 95% humidity. All cell lines were tested and were negative for mycoplasma contamination using a routine mycoplasma PCR kit (ABM).

#### Establishment and Monitoring of Experimental Metastasis

For the single-injection OvCa model, 4×10^5^ ID8-LARRY.GFP or ID8-Luciferase were injected into the peritoneal cavity. To ensure adequate library coverage, 2.4×10^6^ ID8-Cas9.Lib tagged cells were injected for CRISPR screening experiments (>1000x coverage). In the repeated injection model, mice received 4×10^5^ cells per injection, administered six times over the course of 12 days. Mice were typically sacrificed 6 weeks after initial dose of cancer cells. 6×10^5^ E0771-LARRY.GFP or E0771-Luciferase tagged cells were injected into the retro-orbital sinus of mice for one time injection lung metastasis model or 1.5×10^5^ cells in six separate injections for the repeated injection model. Mice were sacrificed between 14-18 days in both experimental models. For monitoring metastasis in luciferase-expressing models, 100 ul of D-luciferin (15 mg/mL stock, GoldBio cat#: LUCK-500) was administered intraperitoneally and imaged using an AMI-HTX spectral imaging instrument. The total emission (flux) data for each region of interest (ROI) was obtained after image analysis using the Aura software.

#### Macrophage Depletion and Histological Analysis

Macrophage depletion was achieved using encapsulated liposomal chlodronate (Encapsula Nano Sciences, SKU# CLD-8901). Mice received a pre-treatment with 200 μl chlodrosome or empty liposome nanoparticles i.p. 24 hours before cancer cell injection. Following pre-treatment, 100 μl of chlodrosome was administered twice weekly throughout the study duration. Omental blocks were collected at endpoint and fixed in 10% neutral buffered formalin (NBF). Fixed tissues were embedded into paraffin blocks and serially sectioned at five non-consecutive intervals separated by 100 μm. Tissue sections were then stained with hematoxylin and eosin (H&E), and images were captured on a Hamamatsu Nanozoomer at the UTSW Whole Brain Microscopy Facility (RRID:SCR_017949) and analyzed in NDPView2 software. For quantification of tumor area, a blinded independent researcher manually segmented metastatic tumor regions and calculated total tumor area for each sample.

#### Individual Gene Knockout Guide Cloning

The CROP-seq-V2 plasmid (Addgene # 127458) was first modified to include a P2A-mCherry sequence downstream of the puromycin resistance gene (PuroR) through a NEBuilder HiFi DNA Assembly (NEB, E5520S) reaction. Then, single-guide RNA (sgRNA) sequences were designed using CRISPick (Mouse GRCm38 reference genome), oligos synthesized by IDT with BsmBI overhangs, annealed, and cloned into the modified CROP-seq-V2 plasmid using the NEBridge Golden Gate Assembly Kit (NEB, E1602S). The resulting plasmids were first transformed into competent *E. coli*, plated on LB-ampicilin (AMP) plates, and grown at 30 °C, before plasmid extraction. Below are the sgRNA sequences used for gene targeting, including the necessary BsmBI overhangs:

sgNTC-1 Top oligo
5’- **CAC CG**A CCT GAT ACG TCG TCG CGT A -3’
sgNTC-1 Bottom oligo
5’- **AAA C**TA CGC GAC GAC GTA TCA GGT **C** -3’
sgNTC-2 Top oligo
5’- **CAC CG**G TAT TAC TGA TAT TGG TGG G -3’
sgNTC-2 Bottom oligo
5’- **AAA C**CC CAC CAA TAT CAG TAA TAC **C** -3’
sgIrf1 guide 1 Top oligo
5’- **CAC CG**T GGA AGC ACG CTG CTA AGC A -3’
sgIrf1 guide 1 Bottom oligo
5’- **AAA C**TG CTT AGC AGC GTG CTT CCA **C** -3’
sgIrf1 guide 2 Top oligo
5’- **CAC CG**C TGT AGG TTA TAC AGA TCA G -3’
sgIrf1 guide 2 Bottom oligo
5’- **AAA C**CT GAT CTG TAT AAC CTA CAG **C** -3’
sgStat1 guide 1 Top oligo
5’- **CAC CG**T ATC CTG TGG TAC AAC ATG C -3’
sgStat1 guide 1 Bottom oligo
5’- **AAA C**GC ATG TTG TAC CAC AGG ATA **C** -3’
sgStat1 guide 2 Top oligo
5’- **CAC CG**G CAA GCG TAA TCT CCA GGT A -3’
sgStat1 guide 2 Bottom oligo
5’- **AAA C**TA CCT GGA GAT TAC GCT TGC **C** -3’
sgIfngr1 guide 1 Top oligo
5’- **CAC CG**G GCT CGG AGA GAT TAC CCG A -3’
sgIfngr1 guide 1 Bottom oligo
5’- **AAA C**TC GGG TAA TCT CTC CGA GCC **C** -3’
sgIfngr1 guide 4 Top oligo
5’- **CAC CG**A CGC ACT CAC CTC CCC ACT C -3’
sgIfngr1 guide 4 Bottom oligo
5’- **AAA C**GA GTG GGG AGG TGA GTG CGT **C** -3’
sgIfngr1 guide 6 Top oligo
5’- **CAC CG**C TGG GCC AGA GTT AAA GCT A - 3’
sgIfngr1 guide 6 Bottom oligo
5’- **AAA C**TA GCT TTA ACT CTG GCC CAG **C** - 3’
sgParp14 guide 1 Top oligo
5’- **CAC CG**A TAG ACT GAG CAT TAA GAC C -3’
sgParp14 guide 1 Bottom oligo
5’- **AAA C**GG TCT TAA TGC TCA GTC TAT **C** -3’
sgParp14 guide 2 Top oligo
5’- **CAC CG**A GGT TCT GAT AAG TAA TGG G -3’
sgParp14 guide 2 Bottom oligo
5’- **AAA C**CC CAT TAC TTA TCA GAA CCT **C** -3’

#### Library Cloning

Six MetTag-LARRY-BC.ID libraries were generated via overhang PCR by amplifying the region within the original pLARRY-eGFP plasmid containing random barcodes. Primers were designed with overhangs containing: (1) a unique 6-nucleotide library identifier (ID), (2) a chromium 10x capture sequence, and (3) BsmBI restriction sites flanking both ends of the barcode (synthesized by Sigma-Aldrich). Amplified barcode fragments were introduced into the pRG212 third-generation lentiviral plasmid using the NEB Golden Gate Assembly Kit (NEB, E1602S) at a 2:1 insert-to-vector molar ratio. The cloning reaction was transformed into Endura electrocompetent *E. coli* cells (Biosearch Technologies, 60242-1) and plated overnight on LB-AMP plates. 3×10^6^-2×10^7^ CFU colonies were harvested by flushing prewarmed LB broth and cultured for 2 hours in liquid LB. sgRNA oligo pools were synthesized as oPools (IDT), containing BsmBI cloning compatible overhangs. The oligos were then annealed and introduced into the CROP-seq-V2.mCherry lentiviral plasmid using the NEB Golden Gate Assembly Kit as described above. Approximately 5×10^7 CFUs for OvCa library and 2×10^7 colonies for MetTag library were harvested for maxiprep. For both LARRY barcode and CRISPR libraries, plasmids were purified using the ZymoPURE II Plasmid Maxiprep kit (Zymo Research, D4202). The MetTag-LARRY-BC.ID vector map and CROP-seq-V2.mCherry vector map will be made available on Addgene upon publication.

#### Plasmid Library Amplification and Diversity Validation

Plasmid DNA was amplified using two step PCR to first amplify and then add indexing sequences to libraries. First round PCR was conducted using custom P5 and P7 adapters with overhangs compatible with Nextera indexing primers, followed by cleanup with Agencourt AMPure XP beads (Beckman Coulter, NC9933872) at a 1.8x bead ratio to remove primers. Index PCR reaction was conducted as outlined in the Illumina 16S Metagenomic Sequencing Library Preparation manual (Illumina, Doc #15044223, page 11-12) using 2x KAPA HiFi HotStart ReadyMix (Roche, KK2601) and Nextera XT Index kit v2 primers (Illumina, FC-131-1001). Final indexed libraries were quantified via the Qubit HS assay kit. LARRY libraries were run on a Bioanalyzer instrument and sequenced on an Illumina MiSeq instrument. CRISPR libraries were run on TapeStation using the High Sensitivity D1000 assay (Agilent, 5067-5585, 5067-5594) and sequenced on a NovaSeq-X instrument. Demultiplexed fastq files were filtered for high quality reads, and the number of unique reads was quantified. For the CRISPR libraries, a customized script was used to estimate the percentage coverage based on the original guide RNA whitelist.

#### Lentivirus Production and Target Cell Transduction

MetTag-LARRY-GFP, CROP-seq-V2.mCherry guide, or Lenti-Luciferase.P2A.Neo (Addgene # 105621), and second-generation lentiviral packaging plasmids were transfected into HEK293T cells along with the CalFectin transfection reagent (SignaGen, SL100478). Lentivirus was harvested after 72 hours and concentrated using Lenti-X Concentrator (Takara Bio, 631231). Concentrated lentivirus was added to target cells along with 8 ug/ml polybrene. To achieve low MOI for CRISPR screening experiments, ID8 p53^-/-^ ovarian cancer cells were infected at a low multiplicity of infection (MOI) of 0.15, as calculated by the percentage of mCherry-positive cells 48 hours post transduction. 48 hours post transfection, cells were selected with either G418 (MetTag-LARRY-GFP lines; Sigma-Aldrich, 4727878001) or puromycin (CROP-seq-V2.mCherry lines; Invivogen, ant-pr-1) for a minimum of 7 days. For clonal and time stamp (BC.ID) tracking, all barcoded pools of cells were infected simultaneously using the same cell passage, same lentivirus titer as measured by a Lenti-X™ GoStix™ Plus kit (Takara Bio, 631280) and were thawed in a staggered manner for injection. This approach ensured equal and short culture time in order to maintain clonal diversity.

#### Genomic DNA (gDNA) Amplicon Library Preparation

Genomic DNA was extracted from cultured cells and digested tissues using either the Quick-DNA Midiprep Kit (Zymo Research, D4075) or the Quick-DNA Miniprep Plus Kit (Zymo Research, D4068), depending on the starting cell number. Extracted DNA yield was quantified via a NanoDrop spectrophotometer, and samples were normalized to the same, lowest input concentration prior to PCR. Two rounds of PCR amplification were performed on > 200 ng starting gDNA material. The first round was utilized for amplicon amplification with nested primers for 15-20 cycles using a high-fidelity hot start polymerase, followed by purification with a 1.8x bead-to-sample AMPure XP bead ratio. Eluted amplicon DNA was quantified using the Qubit high sensitivity (HS) assay kit (Thermo Fisher Scientific, Q33231) and concentrations normalized prior to indexing. Approximately 5 ng of DNA was carried into the index PCR rection, which was conducted similarly to the plasmid library construction protocol described above. Index products were purified using the same AMPure XP beads ratio to remove excess primers. Library quantity was assessed via the Qubit HS assay kit and analyzed on a TapeStation instrument using the High Sensitivity D1000 assay to confirm expected library size and distribution. LARRY gDNA libraries were sequenced in-house on an Illumina MiniSeq instrument (Mid output kit, 300 cycles) while CRISPR gDNA libraries (101 genes*10 guides per gene, 20 nontargeting guides = 1,030 guides) were sequenced on a NovaSeq-X instrument. High quality reads were filtered and analyzed via two parallel approaches: LARRY barcode counts were extracted to quantify library batch identity (ID), while CRISPR libraries were processed using the MAGeCK pipeline to identify enriched or depleted guide RNAs. Primers used for amplification were as follows:

LARRY Round 1 PCR REV, Tm: 55.9:
5’ – GTC TCG TGG GCT CGG AGA TGT GTA TAA GAG ACA GTG GGG AAT TTC TAC TCT TGT AGA TCG – 3’
LARRY Round 1 PCR FWD Stagger, Tm: 59.0:
5’-TCG TCG GCA GCG TCA GAT GTG TAT AAG AGA CAG CTT GCT AGG ACC GGC CTT AAA GC -3’
CRISPR Round 1 PCR REV, Tm: 51.0:
5’ – GTC TCG TGG GCT CGG AGA TGT GTA TAA GAG ACA GTT CTT GGC TTT ATA TAT CTT GTG G– 3’
CRISPR Round 1 PCR FWD Stagger, Tm: 53.0:
5’ – TCG TCG GCA GCG TCA GAT GTG TAT AAG AGA CAG CTC AAG TTG ATA ACG GAC TAG C– 3’

#### PiggyBac Transposon Transfection

SpCas9,CreERT2, and DTR coding sequences were cloned into PiggyBac transposon plasmids containing a hygromycin resistance cassette via Gateway cloning (ThermoFisher Scientific, 11791020 & 11789020). Target cells were plated in 6-well plates 24 hours prior to transfection. SpCas9, CreERT2, or DTR PiggyBac plasmid was co-transfected with Super PiggyBac Transposase (System Biosciences, PB210PA-1), along with the Calfectin transfection reagent. The transfection mixture was added to cells in a dropwise manner. 12-16 hours post transfection complete growth media was replenished. Forty-eight hours post transfection, cells were selected with hygromycin B (Corning, 30-240-CR) for a minimum of 10 days to enrich for successful transgene integrated cells.

#### Cell Preparation for Single Cell RNA Sequencing

Naïve and tumor bearing mice were sacrificed via cervical dislocation six weeks following initial cancer cell injection. To collect peritoneal cells, 5 mL of cold 1xHBSS was injected intraperitoneally using a 25G needle, followed by lavage and peritoneal fluid retrieval using a 20G needle. Omenta were excised from the peritoneal cavity, minced, and digested for 30 minutes at 37°C in RPMI-1640 medium containing 0.4% collagenase type II (Gibco, 17101015), 50 ug/ml DNAseI, and 0.1% BSA. Following digestion, cell suspensions were passed through a 70-uM strainer and washed with cold 1xPBS. Lavage, ascites, and omental samples were resuspended in 1x RBC lysis buffer (BioLegend, 420301) to remove red blood cells. To prepare high viability (>80%) cell suspensions for single-cell analysis, omental samples were stained with ZombieRed (BioLegend, 423109) to mark dead cells and live cells were sorted via FACS. Sorted cells were then prepared for 10X Genomics Chromium Single Cell Gene Expression analysis. For TNBC lung sample scRNA-seq, mice were perfused with 5mL of ice cold 1xPBS, lungs were collected and imaged using the GFP channel on a dissecting scope. Lungs with detectable tumor burden were cut into small pieces and processed into single cell suspensions by digestion with 1x collagenase/hyaluronidase (STEMCELL Technologies, #07912) and 0.15 mg/ml DNase I in serum free RPMI 1640 medium. After passing the digested samples through a 70-uM strainer and washing with serum free RPMI, samples were sorted for GFP+, RBCs were lysed and processed similarly to OvCa samples downstream. A detailed protocol for sample multiplexing via hashtags can be found in the CITE-seq/cell hashing protocol on the CITE-seq website (https://cite-seq.com/wp-content/uploads/2019/02/cite-seq_and_hashing_protocol_190213.pdf). In brief, samples were first blocked with FcR Blocking Reagent (Miltenyi Biotec, 130-092-575) in 50 μl of cell staining buffer for 20 minutes on ice. Samples were then stained with TotalSeq™-A hashing antibodies obtained from BioLegend, including:

M-HTO-1 (M1/42; 30-F11, 155801),
M-HTO-2 (M1/42; 30-F11, 155803),
M-HTO-3 (M1/42; 30-F11, 155805),
M-HTO-4 (M1/42; 30-F11, 155807),
M-HTO-5 (M1/42; 30-F11, 155809),
M-HTO-6 (M1/42; 30-F11, 155811),
M-HTO-7 (M1/42; 30-F11, 155813),
M-HTO-8 (M1/42; 30-F11, 155815).

Staining was performed for 25 minutes on ice, followed by three washes with staining buffer (decreasing EDTA concentration per wash) and loading onto a 10X Genomics Chromium Controller.

#### scRNA-seq Library Preparation and Illumina Sequencing

Single-cell transcriptomic libraries were prepared as outlined in the Chromium Next GEM Single Cell 3’ Reagents Kits v3.1 with Feature Barcoding technology kit manual. Kits necessary for transcriptome library preparation were: Chromium Next GEM Single Cell 3ʹ GEM, Library & Gel Bead Kit v3.1, 4 rxns (10x Genomics, PN-1000128), Chromium Single Cell 3ʹ Feature Barcode Library Kit, 16 rxns (10x Genomics, PN-1000079), Chromium Next GEM Chip G Single Cell Kit, 16 rxns (10x Genomics, PN-1000127), and Single Index Kit T Set A, 96 rxns (10x Genomics, PN-1000213). Hashtag libraries were prepared using the standard CITE-seq protocol (https://cite-seq.com/wp-content/uploads/2019/02/cite-seq_and_hashing_protocol_190213.pdf ). LARRY-BC.ID barcode libraries were constructed using a custom nested PCR protocol followed by indexing. Primers required for LARRY-BC.ID library preparation included:

Round 1 PCR FWD, Tm: 61.0
5’ – *GCA GCG TCA GAT GTG TAT AAG AGA CAG* – 3’
Round 1 PCR REV: 45.0
5’ – CCT TGG CAC CCG AGA ATT CCA aat ttc tac tct tgt aga t – 3’
Index PCR FWD, Tm: 61.0
5’ – AAT GAT ACG GCG ACC ACC GAG ATC TAC ACT CGT CG*G CAG CGT CAG ATG TGT ATA AGA GAC AG* – 3’
Index PCR REV (ADT-RPI-1), Tm 61.0:
5’ – CAA GCA GAA GAC GGC ATA CGA GAT CGT GAT GTG ACT GGA GTT <u>CCT TGG CAC CCG AGA ATT CCA</u> – 3’
*{Partial Read 1N;* LARRY transcript (lowercase), P7 adaptor, RPI1 **index***}*

The quality and quantity of the three individual libraries (transcriptome, HTO, and LARRY) were assessed via Qubit and TapeStation prior to sequencing. Libraries were pooled for sequencing on an Illumina NovaSeq X Plus platform with paired end reads (PE150).

#### scRNA-seq General Data Analysis

Raw fastq files generated from the sequencing runs (mRNA, HTO, LARRY outputs) were processed through the 10x Genomics CellRanger pipeline v.7.2 to generate count matrices. Gene expression (mRNA) reads were aligned to a custom mouse reference genome containing fluorescent marker transcripts (mm10_3.0.0_FPs). Data from two different chromium 10x lanes, ascites lane and omentum lane, were merged using the SCT-normalized dataset integration method. Details related to integration steps can be found at: https://satijalab.org/seurat/articles/. In brief, individual lane single cell data was normalized for HTO (HTOdemux) followed by SCTransform prior to integration. Downstream analyses including dimensional reduction via principal component analysis (PCA), clustering and UMAP embedding, and differential gene expression (DEG) analysis were performed in RStudio under the Seurat package (v4.0)^70^. Cell type identities were assigned based on cluster marker genes. Subcluster level labels indicating different cellular statuses were determined through a combination of cluster marker genes and examining enriched pathways. For each HTO sample, LARRY barcode counts were converted to relative frequencies. This transformation allowed for barcode abundance visualization using the same enrichment scale on a heatmap and for comparisons across biological repeats (HTOs). RNA velocity analysis was performed in Python using scVelo and CellRank. Loom files were generated for each lane and merged, followed by Seurat object conversion to H5AD format. Metadata containing clustering and HTO information was integrated into the SCANPY anndata object. More details on the CellRank method can be found at: https://cellrank.readthedocs.io/en/latest/index.html. NicheNetR and CellChat were performed in RStudio. Sender and receiver cell types were selected prior to NicheNetR analysis. scRNA-seq expression data was analyzed using the Wilcoxon rank sum test either through the direct function wilcox.test() when assigning custom module scores or through the FindMarkers() function as part of the standard DEG analysis. Adjusted p values <0.05 were considered significant.

#### scRNA-seq Pathway Level Analysis

Over-representation analysis (ORA) was performed on a filtered list of upregulated or downregulated genes from DEG analysis and the ReactomePA and clusterProfiler RStudio packages. org.Mm.eg.db package was used to convert genes to Entrez ID symbols. Adjusted p values <0.05 and log fold change > (+/-) 0.25 were considered significant. ORA uses Fisher’s exact test to determine enriched gene sets. For Single Cell Gene Set Enrichment Analysis (ssGSEA), we utilized the escape R package^20^ (https://github.com/BorchLab/escape). In brief, murine Hallmark gene sets were obtained directly within escape using the function getGeneSets (species = “Mus musculus”, library = “H”). Per-cell enrichment scores were calculated with runEscape using the UCell method, with a minimum gene set size of 5, and stored as a separate assay (escape.UCell_H) within the Seurat object. The parameters sets.size = 2000 and sets.n = 5 were used as specified in the script. The resulting enrichment score matrix was used for downstream visualization and clustering (UPAP embedding using Feature Plot). To identify pathways enriched in individual clusters, differential pathway analysis was performed using Seurat FindAllMarkers on the enrichment assay with the Wilcoxon rank-sum test. Only positively enriched pathways were retained for downstream visualization (minimum detection fraction = 0.1, log2 FC threshold = 0.1). Top enriched pathways were ranked by average log2 FC and visualized using violin plots.

#### Flow Cytometry

Samples were extracted and pre-processed similarly as the scRNA-seq samples. Following tissue digestion of lung or omentum tissues and RBC lysis, digested samples were washed in 1x Flow Staining Buffer (Tonbo Biosciences, TNB-4222-L500). For viability assessment, cells were incubated with ZombieAqua Fixable Viability Kit (BioLegend, 423102) at a 1:750 dilution in 1xPBS for 20 minutes at room temperature. Following viability staining, cells were blocked with 1x FcR Blocking Reagent (Miltenyi Biotec, 130-092-575) to minimize nonspecific antibody staining. Cells were then stained with a combination of the following fluorochrome-conjugated surface antibodies: Pacific Blue™ anti-mouse CD45 (BioLegend, 157211, S18009F), APC/Fire™ 750 anti-mouse CD3 (BioLegend, 100247, 17A2) or PE/Cyanine7 anti-mouse CD3 (BioLegend, 100219, 17A2), PE anti-TREM-2 (BioLegend, 824805, 6E9), APC anti-mouse CCR5 (BioLegend, 107011, HM-CCR5), Brilliant Violet 605™ anti-mouse/human CD11b (BioLegend, 101237, M1/70), APC anti-mouse I-A-I-E (BioLegend, 107613, M5/114.15.2), Brilliant Violet 711™ anti-mouse CD4 (BioLegend, 100447, GK1.5), PE/Dazzle™ 594 anti-mouse NK-1.1 (BioLegend, 156517, S17016D) or PE anti-mouse NK-1.1 (BioLegend, 156503, S17016D), Brilliant Violet 421™ anti-mouse PD-L1 (BioLegend, 124315, 10F.9G2), PE/Cyanine7 anti-mouse CD8a (BioLegend, 100721, 53-6.7) or APC/Cyanine7 anti-mouse CD8a (BioLegend, 100713, 53-6.7), Brilliant Violet 785™ anti-mouse F4/80 (BioLegend, 123141, BM8). Cancer cells harvested from culture were similarly stained with ZombieAqua, FcR blocked, and then incubated with IFNGR1 Recombinant Rabbit Monoclonal Antibody FITC (Invitrogen, MA5-46853, 062) with the corresponding Rabbit IgG Isotype Control FITC, eBioscience™ (Invitrogen, 11-4614-80). Cells were then washed with staining buffer, followed by analysis on a Cytek Northern Lights flow cytometer. FCS files were exported and analyzed using FlowJo (BD Biosciences).

#### Western Blotting

Cells were plated at a density of 250,000-500,000 cells per 10 cm dish for regular attachment plate experiments. For anoikis assays, 3x 6 wells per sample were seeded with 300,000 cells each. The next day, cells were either treated with vehicle (1xPBS) or 100 ng/ml mouse recombinant IFNγ (PeproTech, AF-315-05-100UG). After treatment, cells were harvested, washed with 1xPBS, and lysed in M2 lysis buffer (50 nM Tris-HCl, pH 7.4, 150 mM NaCl, 1% Triton X-100, 0.5 mM EDTA) with 1x Protease/Phosphatase Inhibitor Cocktail (CST, 5872). Cell suspensions intended for PARP14 blots were treated with RIPA lysis buffer (Thermo Scientific, Cat#89901) and Pierce Universal Nuclease (Thermo Scientific, Cat#88700). Cells were lysed for 1 hour on ice before centrifugation at high speed to collect protein lysates. Protein quantity was determined using the Pierce^TM^ BCA Protein Assay Kit (Thermo Scientific, 23225). Protein lysate samples utilized for GBP-1 and PARP14 detection were resolved using 4-20% Mini-PROTEAN TGX Precast Protein Gels (Bio-Rad, 4561094). Other immunoblots were performed on fixed concentration using 8-10% SDS-polyacrylamide gels. All gel transfers were performed onto nitrocellulose membranes (Fisher Scientific, 45-004-075). Nitrocellulose membranes were blocked in 5% BSA/1xTBST for 1 hour at room temperature, followed by primary antibody incubation overnight on a 4°C shaker. The following day, blots were washed with 1xTBST followed by incubation with rabbit or mouse-specific horseradish peroxidase (HRP)-conjugated secondary antibodies. Signals were developed with SuperSignal West Pico PLUS Chemiluminescent Substrate (Thermo Fisher Scientific, #34577) or SuperSignal West Femto Maximum Sensitivity Substrate (Thermo Fisher Scientific, #34094). Primary antibodies used for western blotting were:

Beta-Actin (13E5) Rabbit mAb, CST #4970, 1:5,000 dilution
GBP-1 Rabbit Polyclonal Ab, {Thermo, PA5-23509}, 1:1,000 dilution
Cas9 (S. pyogenes) (7A9-3A3) Mouse mAb, CST #14697, 1:1,000 dilution
IRF-1 (D5E4) XP® Rabbit mAb, CST #8478, 1:1,000 dilution
Cre Recombinase (D7L7L) XP® Rabbit mAb, CST #15036, 1:1,000 dilution
GAPDH (14C10) Rabbit mAb, CST #2118, 1:5,000 dilution
Stat1 (D1K9Y) Rabbit mAb, CST #14994, 1:1,000 dilution
p-Stat1 (Tyr701) (58D6) Rabbit mAb, CST #9167, 1:1,000 dilution
PARP-14 (C-1) Mouse mAb, Santa Cruz sc-377150, 1:2,000 dilution
p-Iκbα (Ser32)(14D4) Rabbit mAb, CST #2859, 1:1,000 dilution
Iκbα (L35A5) Mouse mAb (amino-terminal antigen), CST #4814, 1:1,000 dilution
Akt (pan) (C67E7) Rabbit mAb, CST#4691, 1:1,000 dilution
p-Akt (Ser473) (D9E) Rabbit mAb, CST #4060, 1:1,000 dilution
Vinculin (E1E9V) XP® Rabbit mAb, CST #13901, 1:5,000 dilution
Anti-rabbit IgG, HRP-linked Ab, CST #7074, 1:5,000 dilution
Anti-mouse IgG, HRP-linked Ab, CST #7076, 1:5,000 dilution

#### Clinical Bioinformatics

To investigate the prognostic value of genes identified in the present study, a total of 29 genes related to interferon response (25 genes) and NicheNet receptor-ligands (4 genes) were used as a gene pool to construct a prognostic signature. Clinical outcome data, including overall survival (OS) and progression free survival (PFS), as well as related bulk RNA-seq data were retrieved from the curatedOvarianData R package^71^. To develop the prognostic model, genes with a multivariate Cox regression p-value < 0.1 were first prioritized. The remaining candidate genes were then ranked by the absolute value of their Cox regression coefficients in descending order and iteratively added to the gene panel until up to 80% of all candidate genes were included. Cox regression models were subsequently trained on the discovery cohort for OS and PFS separately using the final selected gene set, and performance was validated in an independent cohort. In each cohort, patients were stratified into "low-risk" and "high-risk" groups according to the median linear risk score derived from multivariate Cox regression. Kaplan–Meier curves were used to illustrate survival outcomes between the two risk groups, and log-rank tests were applied to evaluate the prognostic discrimination of the selected gene signature. Patient scRNA-seq RDS file was obtained from Mendeley Data^69^ (https://doi.org/10.17632/rc47y6m9mp.1) and contained 14 patient samples including PBMCs, lymph nodes, primary tumor, metastasis, and malignant ascites. The RNA data was re-normalized and analyzed using the Seurat package in RStudio using a similar approach to the murine scRNA-seq dataset described above.

#### Quantitative Reverse Transcriptase (RT) PCR

The IFNγ treatment time course was performed in 12 well plates. After seeding 1.0×10^5^ cells per well, wells were treated with 100 ng/ml IFNγ for 0 hours (PBS control), 4 hours, 8 hours, or 24 hours. Total cell RNA was extracted from cells using the Direct-zol RNA Miniprep kit with TriReagent (Zymo Research, R2051-A) following the manufacturer protocol. One microgram of total RNA was used for reverse transcription to synthesize single stranded cDNA using the iScript^TM^ Reverse Transcription Supermix for RT-qPCR (Bio-Rad, 1708840). qPCR reactions were performed on a Bio-Rad CFX Opus instrument with SsoAdvanced Universal SYBR Green Supermix (Bio-Rad, 1725270). Data analysis was performed using the CFX Maestro Software. Cq values were used to calculate relative gene expression levels. In brief, calculations were based on the 2^-ΔΔCT^ method with housekeeping genes serving as internal controls and fold change was calculated by normalizing to the zero-hour treatment timepoint Cq values. Statistical analyses and plots were generated in GraphPad Prism. RT-qPCR primers used were as follows:

Beta-actin (Mouse) Forward primer TM: 57.6
5’- GGC TGT ATT CCC CTC CAT CG -3’
Beta-actin (Mouse) Reverse primer TM: 55.9
5’- CCA GTT GGT AAC AAT GCC ATG T-3’
GAPDH (Mouse) Forward primer TM: 59.8
5’- CAT CAC TGC CAC CCA GAA GAC TG -3’
GAPDH (Mouse) Reverse primer TM: 61.6
5’- ATG CCA GTG AGC TTC CCG TTC AG -3’
Gbp2b (Mouse) Forward primer TM: 58.5
5’- ATC TGC TCA TTG CTC AGA CTT ACT GG -3’
Gbp2b (Mouse) Reverse primer TM: 57.4
5’- AAG TAT TTT CTC AGC ATC ACT TCA GCC -3’
Irf1 (Mouse) Forward primer TM: 60.4
5’- ACC AAA TCC CAG GGC TGA TCT GG -3’
Irf1 (Mouse) Reverse primer TM: 58.3
5’- GAA GTT TGC CTT CCA TGT CTT GGG-3’
Stat1 (Mouse) Forward primer TM: 55.1 5’- GCT GCC TAT GAT GTC TCG TTT -3’
Stat1 (Mouse) Reverse primer TM: 55.2 5’- TGC TTT TCC GTA TGT TGT GCT -3’
Marco (Mouse) Forward primer TM: 61.1
5’- ATC CAC CTG ATC CTG CTC ACG G -3’
Marco (Mouse) Reverse primer TM: 60.8
5’- CAC TGC AGC GAG AAG AAG GGC -3’
Slfn1 (Mouse) Forward primer TM: 60.9
5’- AGG GAA AGG TCA AGG CTC ACA TTA AAA ATC C -3’
Slfn1 (Mouse) Reverse primer TM: 60.1
5’- AGG CTT CCC AAC AGA GAC ATC TGG -3’
Parp14 (Mouse) Forward primer TM: 59.8
5’- AAG CCT CAG CCT CTA AGA AGT CAA GG -3’
Parp14 (Mouse) Reverse primer TM: 60.6
5’- CAG GTT TTC AAA TGC CAC CAT GGA GG -3’
St6gal1 (Mouse) Forward primer TM: 58.4
5’- CTG CAG GAT CTC TGA AGA ACT CCC -3’
St6gal1 (Mouse) Reverse primer TM: 59.0
5’- TGT ACA AAC TGT CCT TCA GGA AGC G -3’
Bcl2a1a (Mouse) Forward primer TM: 57.9
5’- CAG ATT GCC CTG GAT GTA TGT GC -3’
Bcl2a1a (Mouse) Reverse primer TM: 59.5
5’- TCT GCA GAA AAG TCA GCC AGC C -3’

#### Anoikis Assays

Cancer cells (1×10^4^) were seeded into 96-well ultra-low attachment (ULA) plates (Corning, CLS7007) to mimic anchorage-independent conditions in the peritoneal cavity environment. After 24 hours, 100 ng/ml recombinant mouse IFNγ or vehicle control (1xPBS) was added to appropriate wells. Twenty-four hours after treatment (i.e. 48hr after start of culture), CellTiter-Glo Luminescent Cell Viability Assay reagent (Promega, G7570) was applied to every assay well, and plates were incubated for 20 min at room temperature. Luminescence, indicating cellular ATP levels, was measured using a Tecan plate reader instrument and analyzed using GraphPad Prism software. For macrophage coculture experiments, 1.0×10^4^ cancer cells were seeded together with 1.0×10^4^ macrophages per well in ULA plates. Macrophages were enriched from the peritoneal cavity of wild type female mice using the MojoSort Mouse F4/80 Positive Selection Kit (BioLegend, 480170), following manufacturer’s protocol. After 48 hours of coculture, cell suspensions were harvested, pelleted, and incubated in TrypLE (Thermo Fisher Scientific, 12605028) for 10 minutes at 37 °C to dissociate cell aggregates. Cell suspensions were passed through a 40-uM strainer before flow cytometric staining as described earlier.

#### Colony Formation Assays

Two hundred ID8 p53^-/-^ cells were plated onto 6 well plates at the start of the experiment. For wells treated with IFN-γ, 100 ng/ml IFN-γ was maintained throughout the 7-day colony formation assay period. Wells were washed with 1xPBS, fixed with methanol, and stained with crystal violet solution (0.5% w/v) at room temperature for 20 minutes. Wells were then washed with deionized water and dried overnight. Per well colony area was obtained in ImageJ and statistical analysis was performed in GraphPad Prism.

#### Cre-based NicheTracing

HEK293T cells were transfected with a loxp-dsRED-loxp-eGFP Cre reporter construct (Addgene # 141148) and cocultured in a 1:1 ratio with Cre.ERT2 expressing ID8 or E0771 cell lines in the presence or absence of 2 uM 4-hydroxytamoxifen (4-OHT, MedChemExpress, Cat # HY-16950). Seventy-two hours after plating, wells were assessed for GFP+ switched cells using a Nikon Ti2 microscope. For independent *in vivo* validation, E0771-Cre.ERT2 cells were injected into the retro-orbital sinus of Ai6 reporter mice. One week post injection, three consecutive tamoxifen or corn oil control treatments were performed (75 mg/kg/dose) i.p.. Two weeks post cancer cell injection, mice were euthanized, perfused intracardially, and lungs were extracted for imaging and flow cytometry analysis. For the ovarian metastasis model, tamoxifen injections were administered in five consecutive doses, two weeks post ID8-Cre.ERT2 cell i.p. injection.

#### Bulk RNA-seq

3.0×10^5^ Ifngr1-KO or sgNTC ID8 p53^-/-^ cells were plated onto 6-well ULA and regular treatment plates. Recombinant IFNγ (100 ng/ml) or vehicle (1xPBS) was added to corresponding wells 24 hours post initial seeding. Cells were harvested 24 hours post IFNγ treatment. ULA samples were subjected to dead cell removal (Miltenyi Biotec, 130-090-101) and total RNA was extracted using Direct-zol RNA Miniprep kit with TriReagent (Zymo Research, R2051-A) following the manufacturer protocol. Purified RNA was subjected to library preparation. In brief, mRNA was purified using poly-T oligo magnetic beads, fragmented, and reverse transcribed. Resulting cDNA was subjected to amplification and indexing. Final libraries were sequenced on a NovaSeq-X instrument. Raw reads were trimmed and aligned to the mouse reference genome (mm10). Following transcript level quantification, subsequent count matrices were analyzed in RStudio. DESeq2 was used for downstream analysis and variance-stabilized counts (VSTs) were used for PCA and visualization. Pathway analysis (Hallmark, Reactome, Go Biological Processes) was conducted through the clusterProfiler R package.

#### Diphtheria Toxin-mediated DTC Ablation

DTR or vector (stuffer) control MetTag-LARRY-GFP cells were injected i.p. in a staggered manner into female C57BL/6J mice according to the timeline established in Figure 1. Two hours post first wave of DTR-expressing cancer cell injection, mice received the first dose of either DT (1 ug DT/100ul PBS; Sigma, D0564) or PBS vehicle i.p. to abolish DTCs before colonizing. DT administration was continued every other day for the duration of cancer cell injection. Animals were monitored daily for DT-induced side effects and sacrificed only two days post final cancer cell injection to limit the duration of DT exposure.

#### Light Sheet Fluorescent Microscopy of Cleared Omental Tissue

4×10^5^ ID8 p53^-/-^ cancer cells expressing an RFP tag or vehicle (1xPBS) were injected intraperitoneally into C57BL/6 background mice. 48 hours later, both groups received a second dose of 4×10^5^ ID8 p53^-/-^ cells labeled with GFP. Omenta were harvested 24 hours post ID8 p53^-/-^ GFP injection. Tissues were surgically harvested, fixed overnight in 4% paraformaldehyde (PFA) at 4 °C with gentle rotation. Tissues were then washed in 1xPBS containing 0.02% sodium azide to remove residual fixative. After fixation, specimens were decolorized overnight with 25% Quadrol at 37°C and then permeabilized overnight with blocking buffer (0.5%NP40,10% DMSO, 5% donkey serum, 0.5% Triton X-100 in 1xPBS) at RT, before immunolabeling. Omental samples were incubated with primary antibodies; RFP preabsorbed antibody (Rockland Immunochemicals INC., 600-401-379) and anti-GFP antibody (Aves Labs, AB_2307313) under gentle rotation for 5 days at RT, washed with wash buffer (0.5%NP40, 10% DMSO in 1xPBS) for 6 hours, refreshing every 2 hours. To reduce fluorescent aggregate formation, samples were post-fixed with 4% PFA and labeled with secondary antibodies: donkey anti-chicken AF488 (Invitrogen, a78948) and donkey anti-rabbit CF555 (Biotium, 20038) for 4 days at RT with gentle rotation. Secondary antibodies were removed with wash buffer for 6 hours, refreshing every 2 hours and refreshing next day. After immunolabeling, tissues were incubated in SYTOX deep red (Invitrogen, S11381) overnight for DNA staining. Following immunostaining, tissues were dehydrated through a graded methanol series (25%, 50%, 75% and 2 x 100%). Samples were then delipidated with dichloromethane until samples sank into the tube. At the end, tissues were cleared by refractive index matching in a benzyl alcohol/benzyl benzoate solution (1:2). This solvent-based clearing workflow reduced light scattering and enabled volumetric imaging of intact or minimally sectioned specimens. Tissues were imaged on a multiscale cleared tissue axially swept light-sheet microscopy (MCT-ASLM) platform. The datasets analyzed were acquired using the high-magnification nanoscale configuration in the 38x imaging format with a pixel size of 0.167 x 0.167 x 0.200 microns. Three-dimensional (3D) fluorescence image stacks were processed to reduce punctate aggregate-like signal before downstream visualization. For each volume, a smooth background estimate was generated with anisotropic Gaussian filtering, and a background-subtracted response image was used to emphasize small, bright structures relative to the surrounding tissue signal. Candidate aggregates were detected from the extreme upper tail of this response distribution using robust statistics, together with an intensity-to-background ratio criterion, so that unusually bright objects could be identified even in heterogeneous fields. Detected seeds were grouped into connected 3D components and expanded locally to capture the immediate aggregate neighborhood while limiting growth into adjacent tissue. Masked regions were then replaced with a local intensity estimate derived from the surrounding image and lightly smoothed to reduce abrupt transitions. Aggregate detection and replacement were repeated iteratively until the residual burden of high-intensity outliers was substantially reduced or no longer detected by the same decision rules. In addition to the cleaned image, the workflow records a cumulative aggregate mask, per-iteration summaries, and preview renderings to facilitate quality control. 3D fluorescence data were visualized interactively in Napari. Individual fluorescence channels were displayed as separate image layers with additive blending and attenuated maximum-intensity rendering to support inspection of spatial overlap in 3D. Channel identities were rendered with consistent lookup tables, while contrast windows were defined on a per-dataset basis to account for sample and region-dependent fluctuations in brightness. Voxel spacing was applied during rendering to preserve the native anisotropy of the image volumes.

## Data and Resource Availability

scRNA-seq datasets with LARRY barcode and HTO multiplexing was deposited at GEO: GSE302619 (ovarian) and GSE309120 (lung) and will be made publicly available upon publishing. Human scRNA-seq data has been published elsewhere and is publicly available: GSE14764. Code and any additional information required to reanalyze the data reported here can be requested from the lead contact.

## Declaration of generative AI and AI-assisted technologies in the writing process

During the preparation of this work the author(s) used AI technologies to improve writing clarity. After using this tool/service, the author(s) reviewed and edited the content as needed and take(s) full responsibility for the content of the publication.

