## Supplemental figures S1-S9 for "Tumor-Intrinsic IFNγ Signaling and Niche Adaptation Drive Early Colonization in Ovarian Cancer Metastasis"

**Supplementary Figure 1:** MetTag Tracing Construct and Metastatic Model Validation

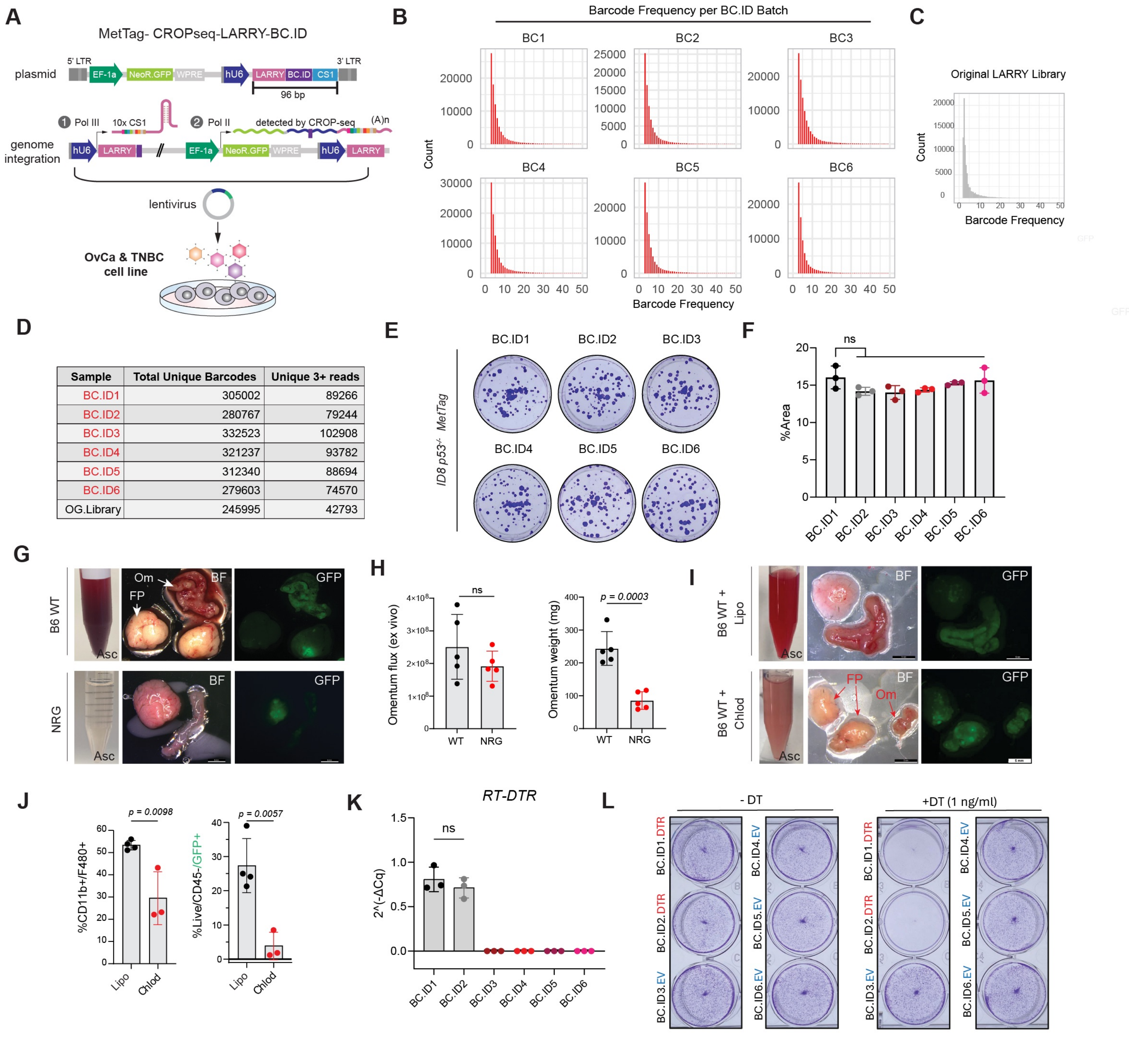


| Sample | Total Unique Barcodes | Unique 3+ reads |
|---|---|---|
| BC.ID1 | 305002 | 89266 |
| BC.ID2 | 280767 | 79244 |
| BC.ID3 | 332523 | 102908 |
| BC.ID4 | 321237 | 93782 |
| BC.ID5 | 312340 | 88694 |
| BC.ID6 | 279603 | 74570 |
| OG.Library | 245995 | 42793 |

**Figure S1. MetTag Tracing Construct and Metastatic Model Validation**
(A) Schematic depicting MetTag library barcoding plasmid, and modes of detection upon plasmid integration: DNA level, Pol III U6 driven RNA level, and Pol II CROP-seq based. (B-C) Barcode frequencies comparing relative evenness of library distribution across BC.ID batches (B) and original depositor LARRY library (C). (D) Table depicting unique reads detected in each library batch after MiSeq sequencing and read filtering. (E) Representative colony formation assay wells for each MetTag-LARRY-BC.ID ID8 p53-/- line. (F) Quantification of per well colony area (n = 3 per sample). Colony areas were compared in GraphPad Prism and plotted as mean with SD and differences between groups were compared via a two-tailed unpaired Student's t test. (G) Representative ascites images and dissecting scope images of GFP+ metastases formed under the repeated injection schema in wild type (n = 4) and NRG mice (n = 4). (H) Separate cohort of mice (n = 5 per group) were subjected to ID8 p53-/- Luciferase injection and omental burden was quantified via ex vivo BLI imaging (left) and by measuring omental weights (right). (I) Representative ascites images and dissecting scope images of GFP+ metastases formed under the repeated injection schema in liposome (n = 4) and chlodro-some treated mice (n = 3). (J) Bar chart depicting macrophage depletion efficiency (left) and quantification of GFP+ cells (right) in ascites under macrophage depletion. (K) RT-qPCR based assessment of DTR transcript expression in each MetTag-LARRY-BC.ID ID8 p53-/- line after trans-fection with DTR or stuffer constructs. Flow cytometry and qPCR data was analyzed in GraphPad Prism and plotted as mean with SD and differences between groups were compared via a two-tailed unpaired Student's t test. (L) Representative colony formation assay wells for each MetTag-LARRY-BC.ID ID8 p53-/- line without (left) or with DT treatment (right).

**Supplementary Figure 2:**
scRNA-seq-based Tumor Cell Identification, *In Vitro* BC Distribution, and Cancer Cell Velocity

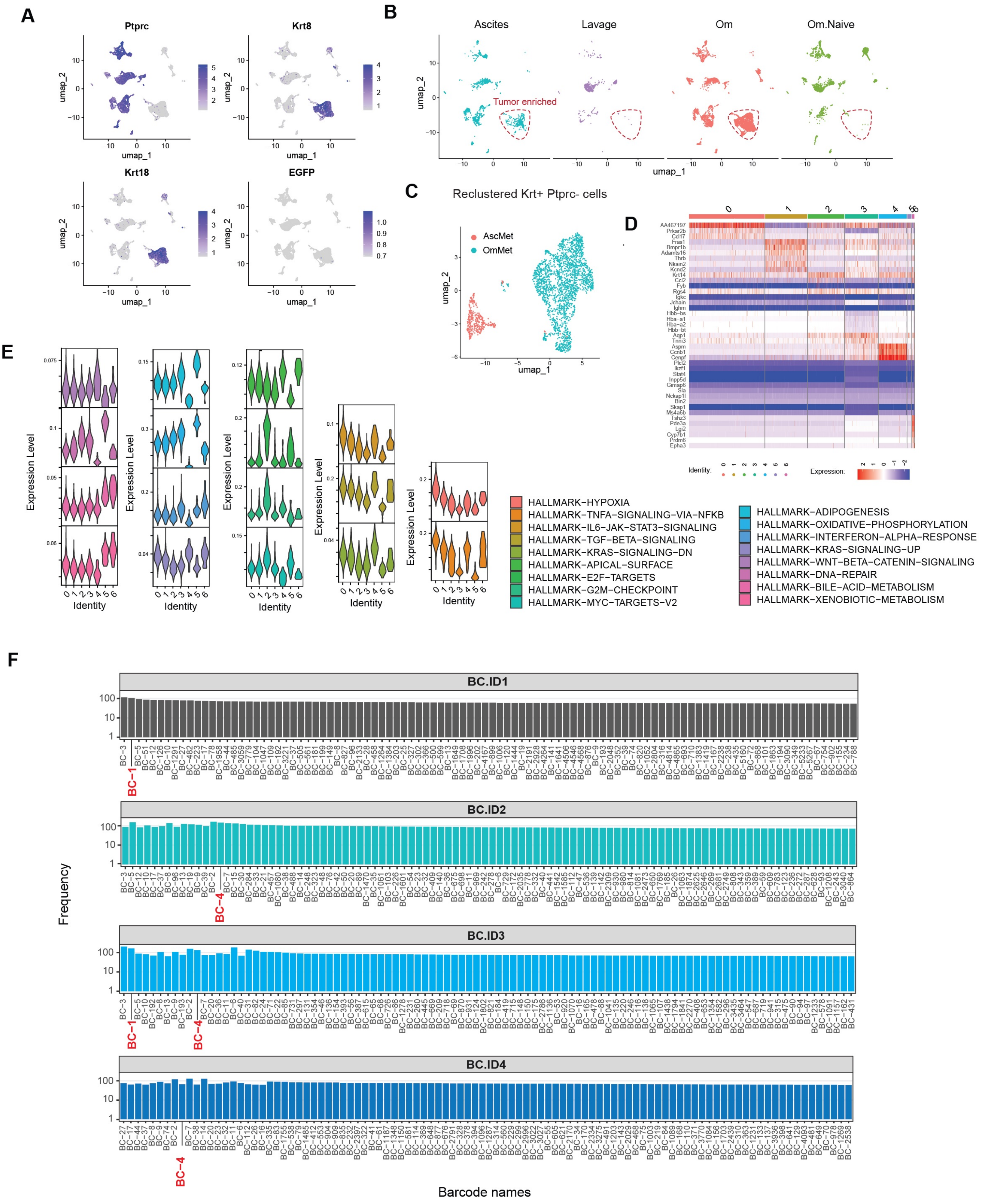

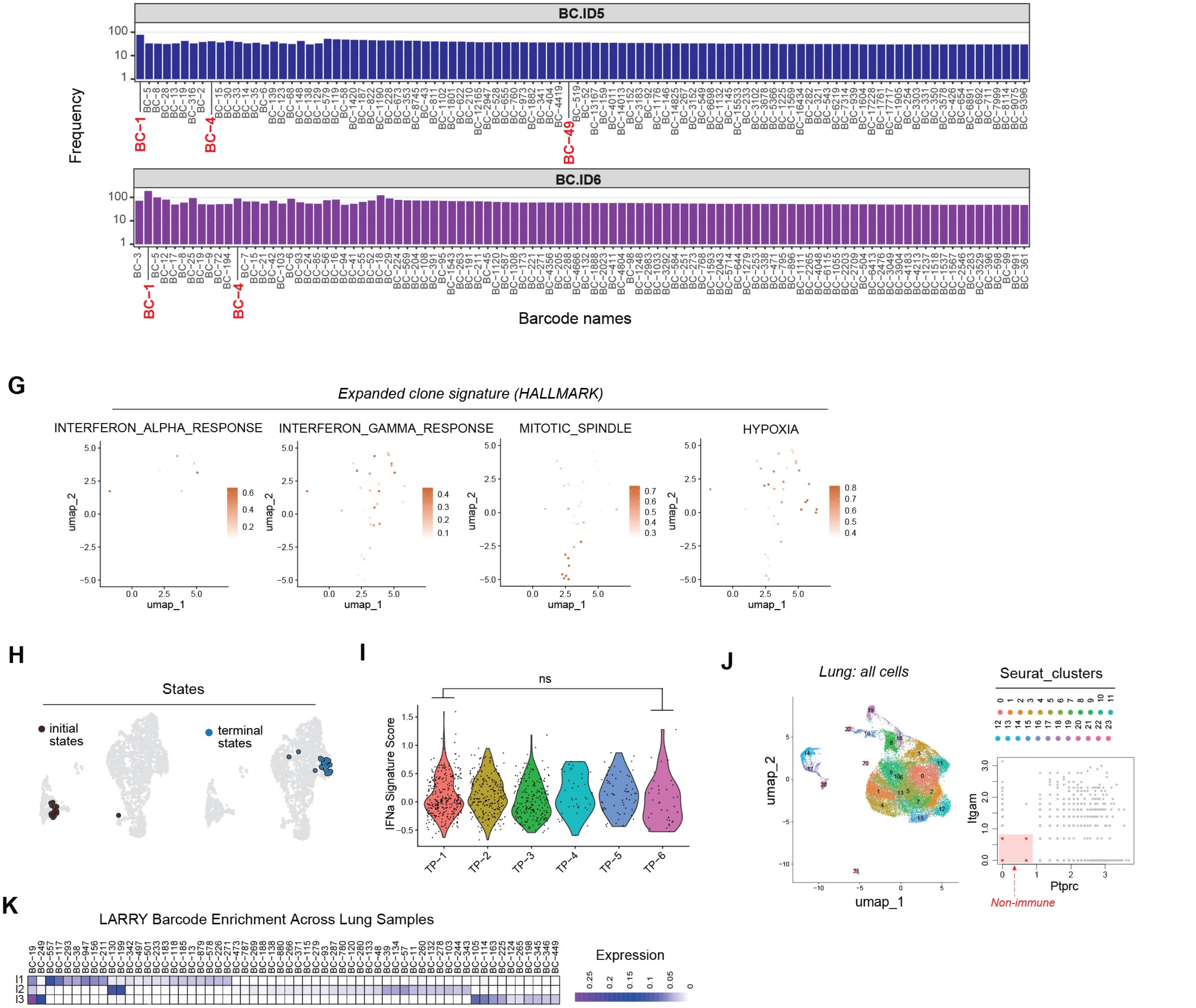


**Figure S2. MetTag Tracing Construct and Metastatic Model Validation**

(A) Feature plot of main marker genes used to define and subset cancer cells. (B) Split UMAP depicting cluster distributions across the four experimental conditions and identifying cancer cell-enriched clusters in ascites and omentum. (C) Reclustered cancer cell UMAP colored by the two metastasis-bearing anatomical locations. (D) Heatmap of reclustered cancer cell marker genes. (E) Violin plots of escape R generated Hallmark pathways each represented by a different color. Details on escape R ssGSEA can be found in the Methods section. (F) Log10-transformed frequency distributions of top 100 LARRY barcodes with counts >= 10, obtained from in vitro cell culture. (G) UMAPs depicting main Hallmark pathway signatures represented in the top three clonally expanded barcoded cells. (H) CellRank-generated initial and terminal cellular states overlayed on the cancer cell UMAP. (I) Violin plot of IFN$\alpha$ signature score across the six different BC.ID timepoints. IFN$\alpha$ signature was assessed by first defining the signature using Hallmark IFN$\alpha$ response genes, followed by directly applying the wilcox.test() function in Seurat to compare expression level differences between TP-1 and TP-6 groups. (J) UMAP of captured single cells in TNBC lung metastasis model (left) and RNA-based gating strategy to subset non-immune cells for subsequent barcode analysis. (K) Heatmap depicting enriched clonal LARRY barcodes in each lung sample (HTO n's = 3).

**Supplementary Figure 3:** IFN Signaling in Metastatic Niches and *In vivo* CRISPR Screening

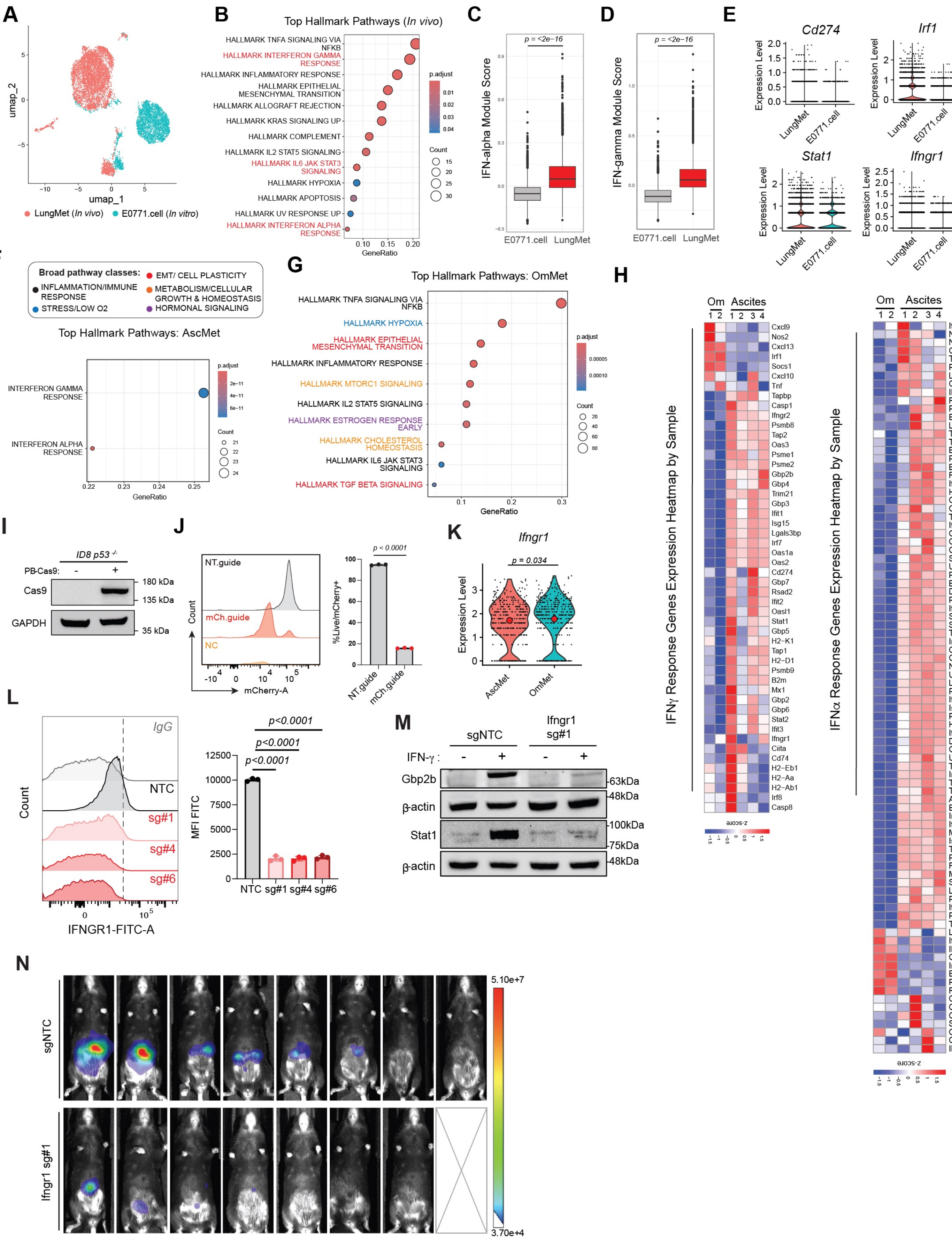

**Figure S3. IFN Signaling in Metastatic Niches and *In vivo* CRISPR Screening**
(A) UMAP of captured cancer cells colored by in vivo (Lung.Met; HTO n's = 2) and in vitro (E0771.cell; HTO n's = 1) cell line. (B) ORA analysis-derived Hallmark pathways enriched in vivo compared to in vitro. Module scores comparing cannonical IFNγ (C) and IFNα (D) genes between the two conditions. (E) VlnPlots of select individual IFNγ response related genes. (F-G) ORA analysis-derived pathways based on DEGs between AscMet and OmMet. (H) Heatmaps depicting individual IFN response gene scoring differences between AscMet and OmMet. (I) Western blot validation of SpCas9 expression in ID8 p53-/- cells after PiggyBac transposon transfection and selection with hygromycin. (J) Representative flow cytometry generated histogram of mCherry fluorescence (left) and bar chart depicting percent mCherry positive cells five days after lenti-guide transduction (right). (K) scRNA-seq derived violin plot of Ifngr1 expression levels in cancer cells in vivo. Module scores and individual gene transcript levels were compared using the wilcox.test() function in Seurat. (L) Representative flow cytometry histogram of IFNGR1 surface expression in knockout vs. wild type cells (left) and bar chart with quantification of IFNGR1 mean fluorescent signal changes (right). n = 3 biological replicates per group were analyzed using a two-tailed Student's t-test and graphed as mean with SD. (M) Western blot probing for IFNγ downstream transcriptional targets, Stat1 and Gbp2b, in Ifingr1 (sg#1) knockout cells. (N) Endpoint BLI flux images of sgIfngr1 (n = 7) compared to sgNTC experiment mice (n = 8).

**Supplementary Figure 4:** IFNγ Downstream Mediators of Anoikis Resistance

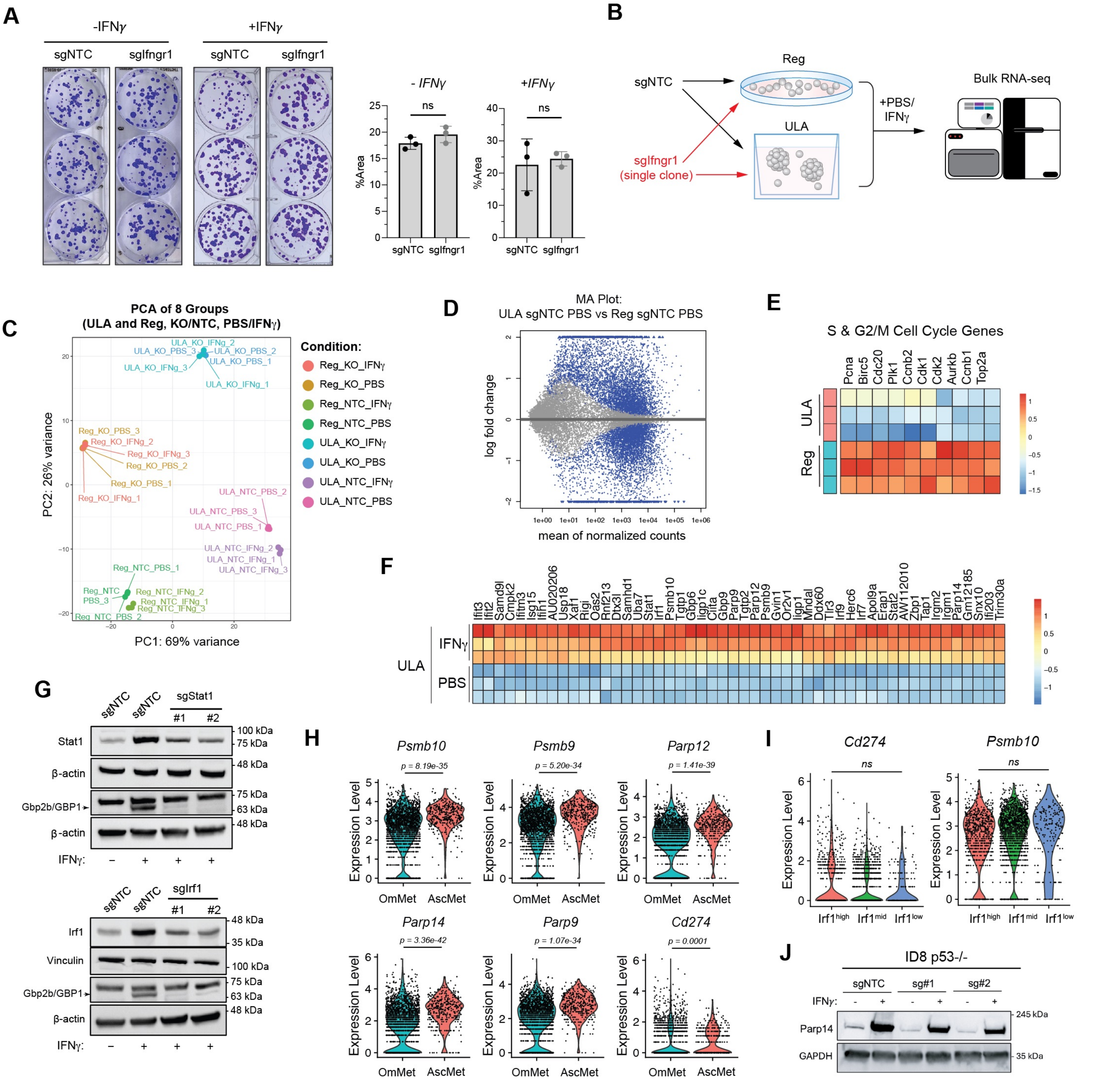

**Figure S4. IFNγ Downstream Mediators of Anoikis Resistance**
(A) Colony area-based quantification of clonogenic assays in absence and presence of recombinant IFNγ, with or without Ifngr1 (sg#1) knockout. Colony area was plotted in GraphPad Prism as mean with SD and analyzed using a two-tailed Student's t test. (B) Schematic depicting in vitro bulk RNA-seq experiment comparing ULA and regular culture conditions under IFNγ treatment. (C) PCA plot of sequenced RNA-seq groups, generated from variance-stabilized transformed (VST) counts. Each dot represents a different sample; color represents different treatment or culture condition. (D) MA quality control plot showing DEGs between ULA and regular culture. Blue points indicate significant genes (padj < 0.05, log2FC>+/- 1). (E) S and G2/M cell cycle gene heatmap comparing wild type sgNTC cells cultured under ULA and regular culture conditions. (F) Heatmap of top DEGs - both unique and overlapped with regular culture - between PBS and IFNγ treated cells under ULA culture (n = 3 per condition). (G) Western blots showing partial knockout of Stat1 and Irf1 transcription factors, as well as subsequent ablation of Gbp2b expression with IFNγ treatment, as a proxy for downstream signaling ablation. (H) Violin plots comparing expression levels of immunoproteasome (Psmb10, Psmb9), DNA repair/apoptosis (Parp12, Parp14, Parp9), and immune checkpoint (Cd274) levels between OmMet (n = 2) and AscMet (n = 4). (I) Violin plots comparing expression level of Cd274 (left) and Psmb10 (right) between Irf1$^{high}$ and Irf1$^{low}$ cells. scRNA-seq based transcript expression levels depicted in (H) and (I) were compared by applying the wilcox.test() function in Seurat. (J) Western blot depicting partial knockout of Parp14 in ID8 p53-/- cells.

**Supplementary Figure 5:** scRNA-seq Analysis of Myeloid and Lymphoid Populations

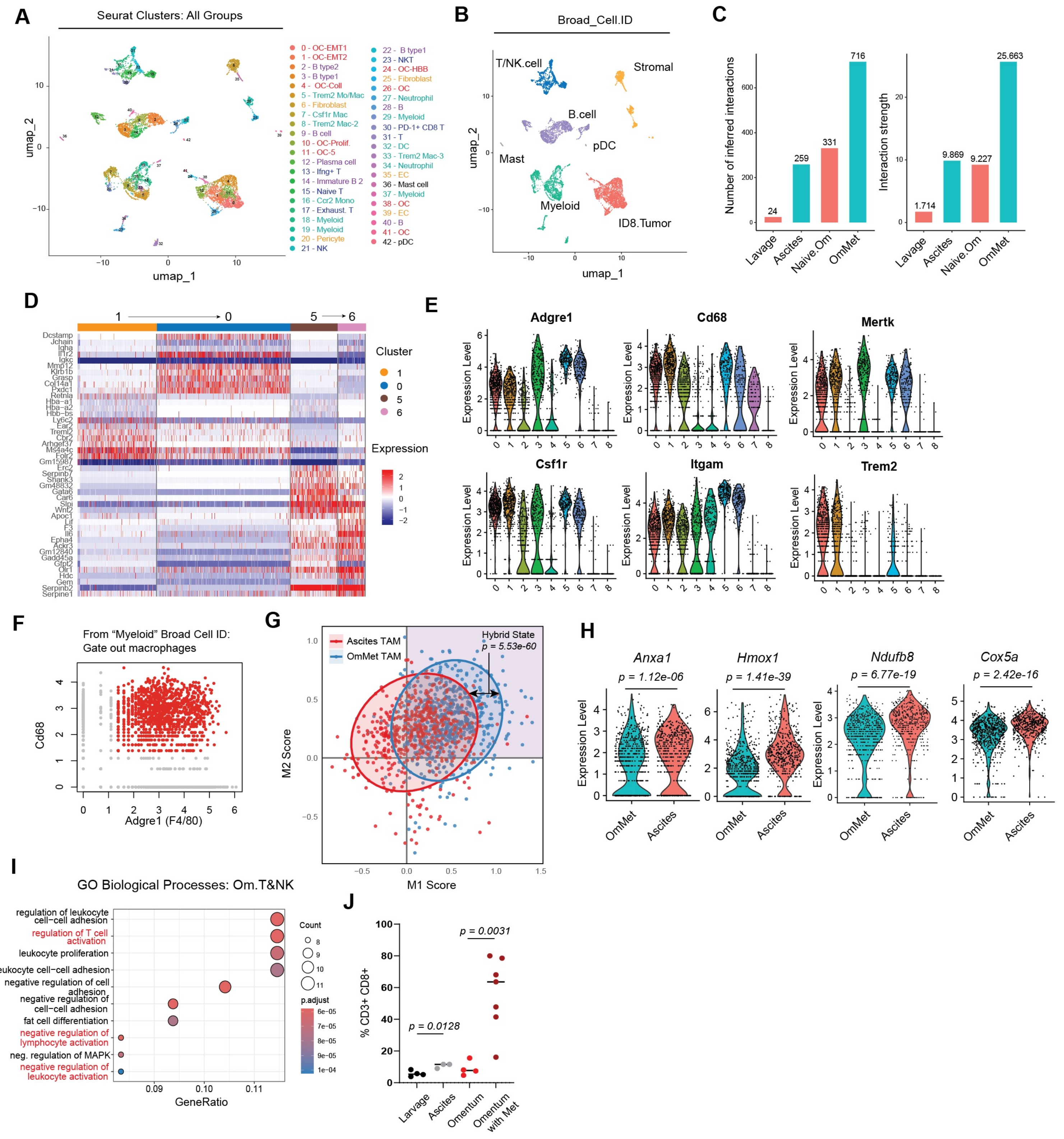

**Figure S5. scRNA-seq Analysis of Myeloid and Lymphoid Populations**
(A) UMAP projection of all cells captured across four sequenced groups, consisting of two integrated datasets. (B) "Broad_cell.ID" classification of Seurat clusters into cell types based on cluster marker genes. (C) CellChat generated bar charts depicting number of interactions (left) and interaction strength (right) between different cell types across the four sample groups. (D) Heatmap of myeloid TAM subclusters 1, 0, 5, and 6. (E) Violin plots showing gene expression of canonical TAM markers across myeloid subclusters. (F) RNA-based gating strategy to subset out macrophages for subsequent analysis. (G) Biaxial scatter plot of M1 and M2 module scores with overlapped ascites and omental TAMs. (H) Violin plots comparing individual genes belonging to cellular response to stress (*Anxa1, Hmox1*) and cellular respiration (*Ndufb8, Cox5a*) pathways between omental and ascites TAMs. Hybrid score depicted in (I) and violin plot expression levels in (H) were compared using the wilcox.test() function in Seurat. (I) Top ORA-derived GO Biological Process pathways upregulated in omentum-derived compared to ascites-derived T/NK cells. (J) Flow cytometry analysis of CD8+ T cells frequencies in naïve and metastatic settings. Flow cytometry data was analyzed via two-tailed unpaired Student's t test and plotted as individual values with median.

**Supplementary Figure 6:** Flow Cytometry Gating Strategies

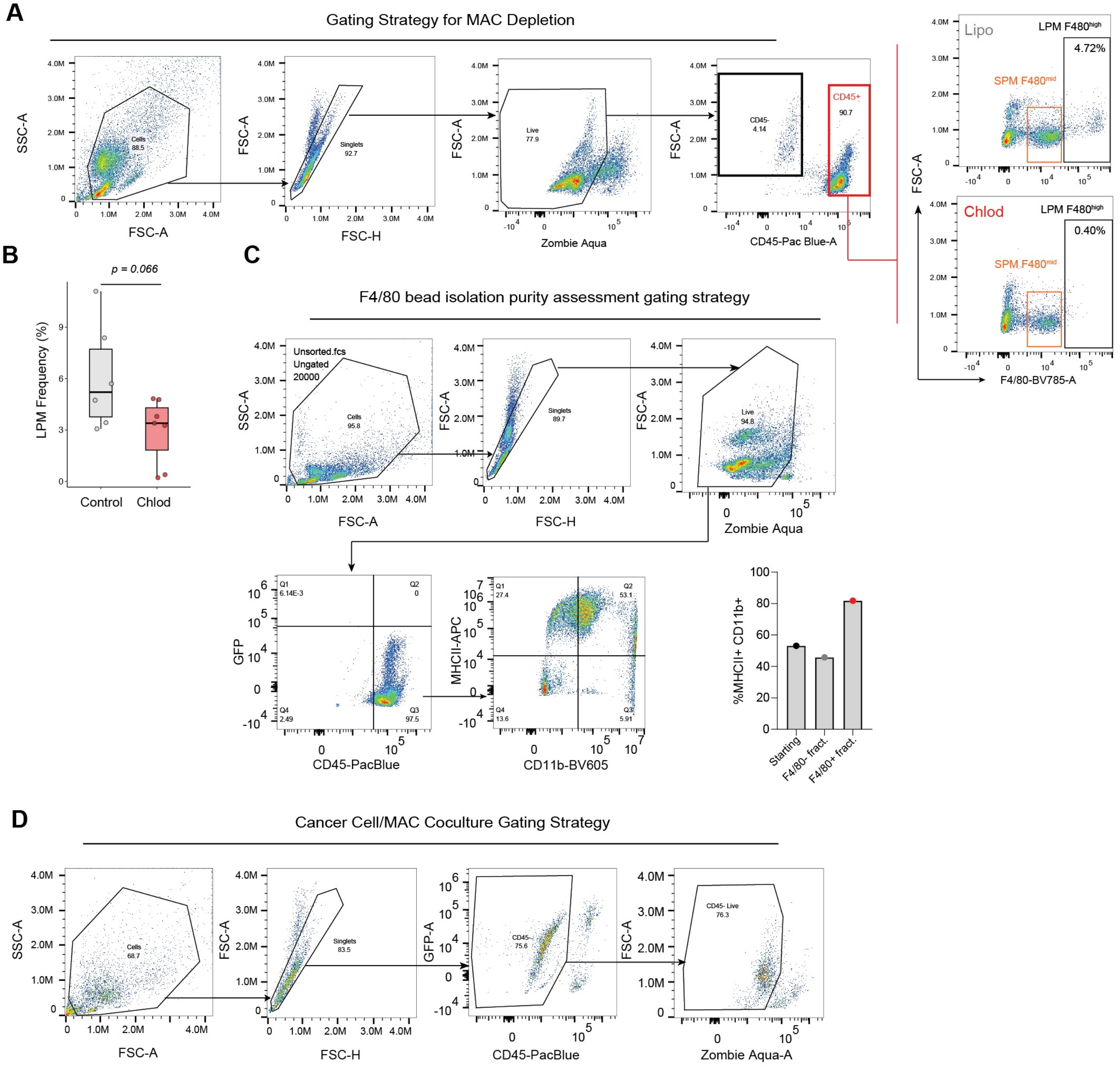


**Figure S6. Flow Cytometry Gating Strategies**

(A) Flow gating strategy for macrophage depletion study and representative gates for F4/80 macrophages in liposome control and chlodrosome treated groups. (B) Quantification of F4/80$^{high}$ LPMs in control and chlodrosome treated mice. P value was generated using a two-tailed Welch's t-test and corresponded to a 51% decrease in macrophage frequency. (C) Flow gating strategy for assessing MHC-II+/CD11b+ macrophage enrichment following F4/80 magnetic bead isolation from peritoneal cavity. Bar plot showing relative enrichment of macrophage populations before the start of coculture. (D) Flow gating strategy assessing CD45- cancer cell viability (Zombie Aqua negative population) after co-culture completion.

### **Supplementary Figure 7:** Mechanistic Exploration of TAM Pro-metastatic Function

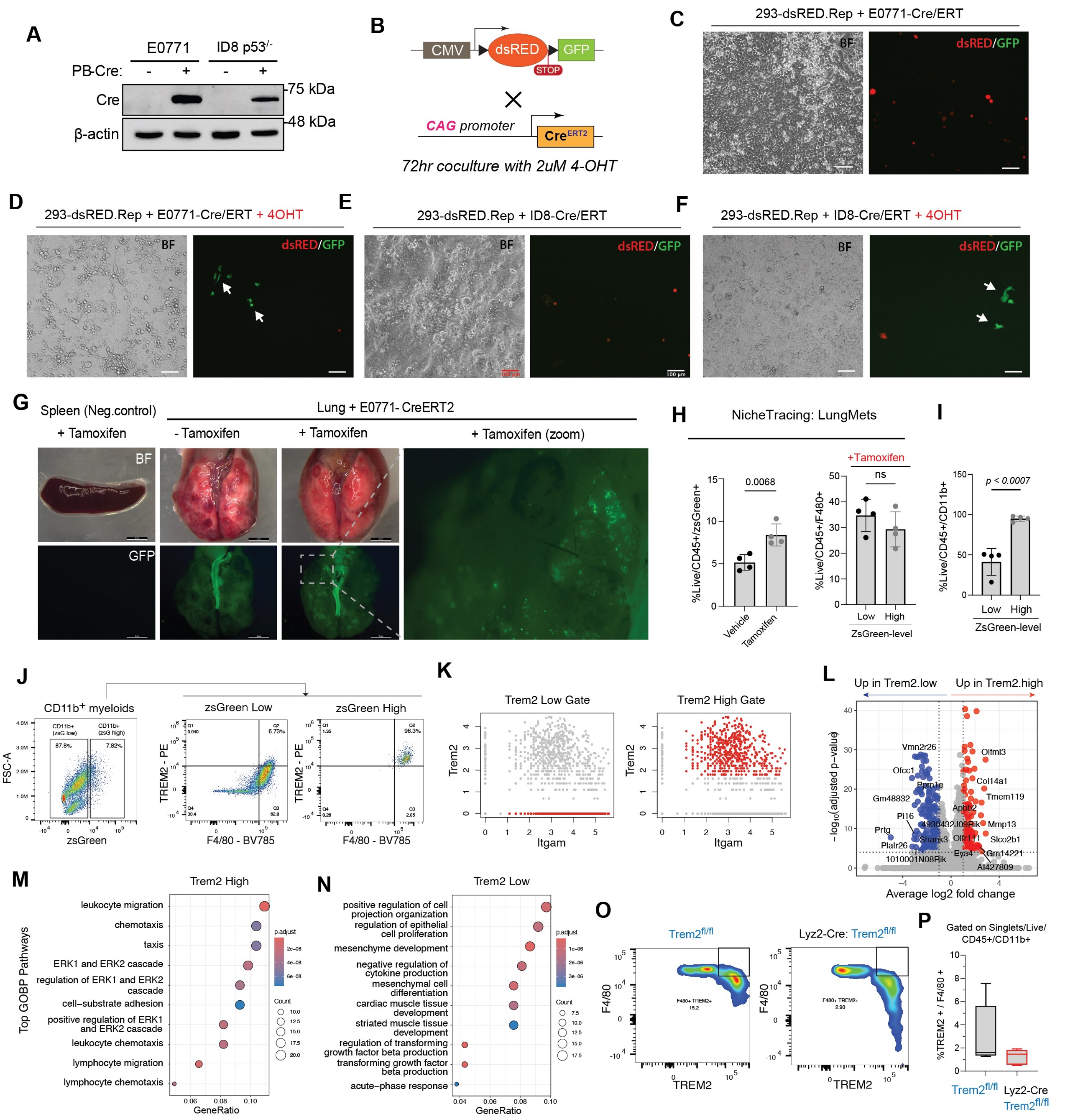


**Figure S7. Mechanistic Exploration of TAM Pro-metastatic Function**

(A) Western blot validation of Cre expression in target cell lines: E0771 murine TNBC line and ID8 p53-/- ovarian cancer cell line. (B) Schematic depicting Cre reporter used to transfect HEK293T cells and Cre constructs expressed by target cells for in vitro color switch assays. (C-F) Fluorescence cell culture images generated after 72 hours of cancer cell-CreERT.2/HEK293T-reporter coculture with or without 2 uM 4-OHT treatment. (G) Representative dissecting microscope images of negative control spleen and lugs collected 2 weeks post E0771-CreERT.2 retro-orbital injection. (H) Flow cytometry bar plots at experimental endpoint showing percent switched immune cells (ZsG+) with (n = 4) or without (n = 4) tamoxifen treatment (left) and within the tamoxifen treated group, frequency of ZsG+ macrophages compared to ZsG- (right) in metastatic lungs. (I) Bar plot showing frequency of ZsG+ CD11b myeloids compared to ZsG- in malignant ascites (n = 4). Flow cytometry data in (H) and (I) was analyzed using a two-tailed unpaired Student's t test and plotted as mean with SD. (J) Representative flow cytometry gating for TREM2 macrophage enrichment in ZsG+ fraction. (K) scRNA-seq gating strategy used to subset TREM2-high and low TAMs. (L) Volcano plot comparing DEGs between metastasis-infiltrating Trem2-low and Trem2-high macrophages. Significant DEGs were generated using the FindMarkers() function in Seurat and Wilcoxon test to compare gated Trem2-low and Trem2-high cells. (M-N) Go Biological Processes pathways upregulated in Trem2-high (left) and Trem2-low (right) TAMs. Macrophage-specific Trem2 depletion validation representative flow cytometry plots (O) and quantification (P).

**Supplementary Figure 8:** Patient and Murine Tumor-TAM NicheNet Analysis and Mechanism

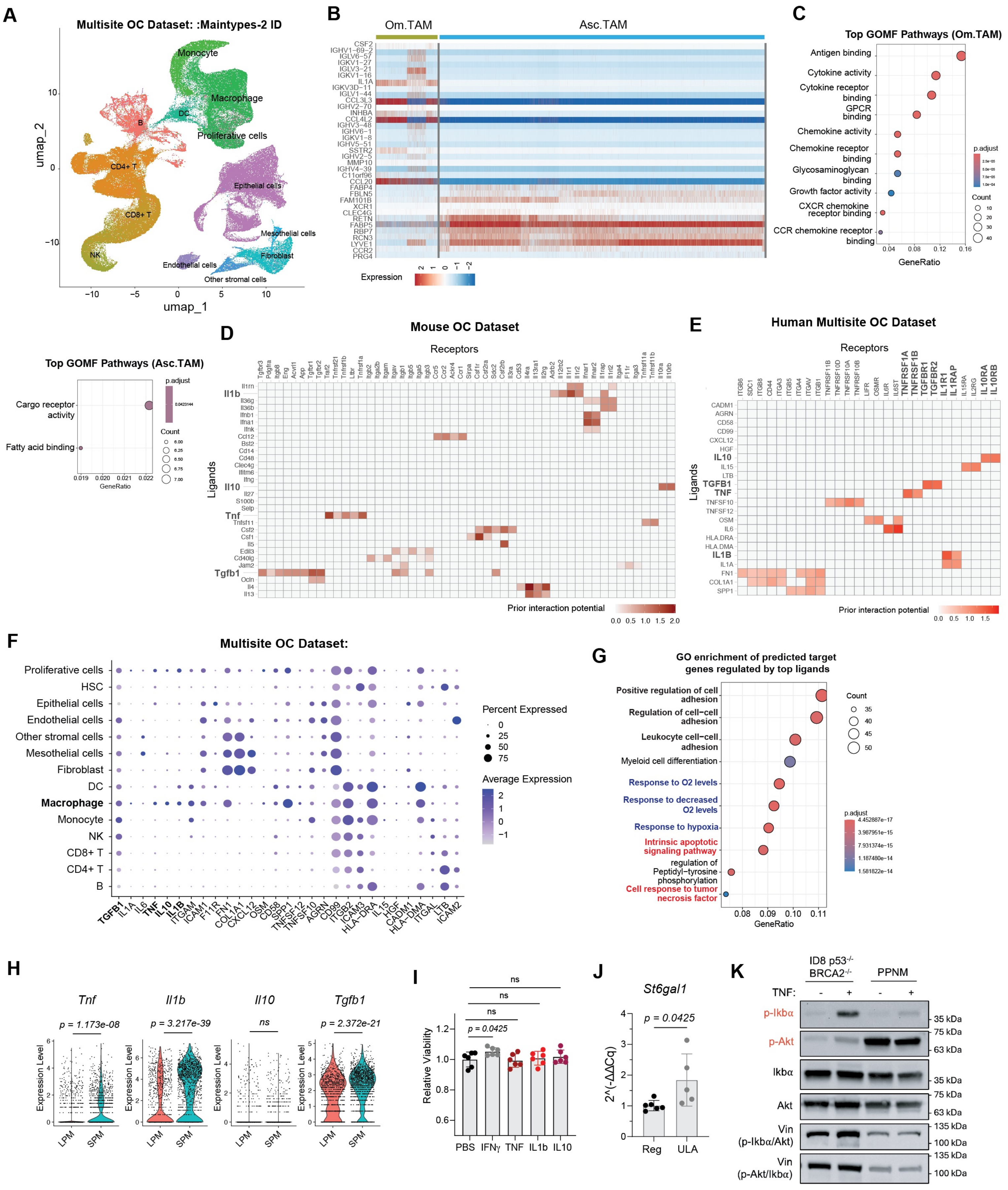

**Figure S8. Patient and Murine Tumor-TAM NicheNet Analysis and Mechanism**
(A) UMAP depicting all (>200,000) single cells collected from five anatomical locations and annotated by cell type ("Maintypes-2"). (B) Heatmap of DEGs between Ascites and Met.Ome-derived macrophages. DEG analysis was performed using the FindMarkers() function in Seurat. (C) Dot plots depicting top GO molecular function pathways expressed in patient omental TAMs (left) and ascites TAMs (right). (D) NicheNet heatmap depicting top receptor-ligands pairs in murine ascites. (E) NicheNet heatmap depicting top receptor-ligands pairs in patient ascites. (F) Bubble plot of NicheNet-derived ligand expression among Maintypes-2 clusters. (G) Dot plot of GO pathways enriched in cancer cells based on regulatory potential and downstream genes affected by macrophage ligands. (H) Murine scRNA-seq violin plots showing transcript levels of *Tnf, Il1b, Il10*, and *Tgfb1* in SPMs and LPMs. Expression levels were compared using the wilcox.test() function in Seurat. (I) Cell-Titer Glo (CTG) assay luminescence assay graphed as fold change for Reg culture, with and without cytokine treatment, and comparing viability to the PBS-treated group. Fold change for each biological replicate (n = 6) was determined by normalizing the cell line's CTG value to the regular attachment, without cytokine treatment, baseline. CTG assay luminescence data was analyzed using a two-tailed Student's t test and graphed as mean with SD. (J) RT-qPCR derived St6gal1 transcript levels in ID8 p53-/- cells under Reg and ULA culture conditions, n = 6 per group where each dot represents the average of technical replicates. Fold change (2^(-ddCq)) data was analyzed using a two-tailed Student's t test and graphed as mean with SD. (K) Western blot probing for p-Ikb$\alpha$/Ikb$\alpha$ and p-Akt/Akt protein levels in ID8 p53-/- BRCA2-/-

### Supplementary Figure 9:

Prognostic Value of Interferon and NicheNet-based Gene Signatures in Ovarian Cancer Patients

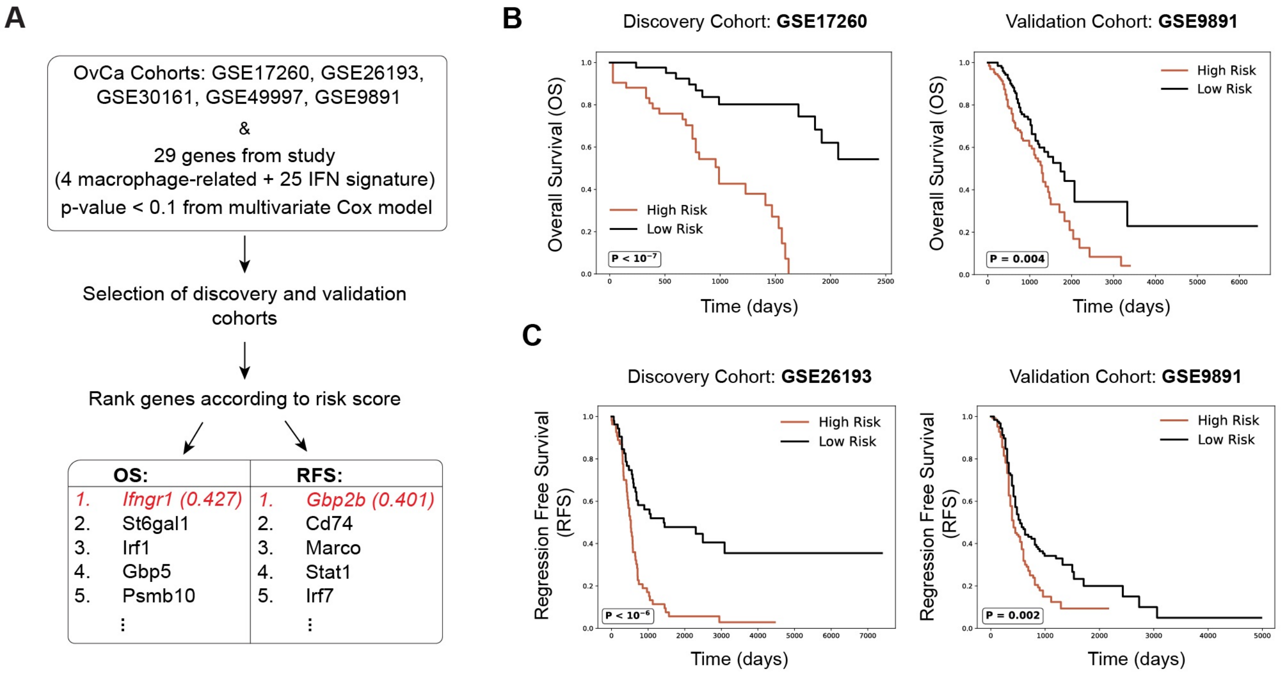


**Figure S9. Prognostic Value of Interferon and NicheNet-based Gene Signatures in Ovarian Cancer Patients**
(A) Bioinformatics workflow using clinical RNA-seq datasets derived from the curatedOvarianData package and 29 genes from the present study (25 IFNγ-related and four NicheNet-derived genes). Genes were prioritized via multivariate Cox proportional hazard regression and iteratively selected from the datasets. (B) Kaplan-Meier plots depicting overall survival (OS) for patients stratified into high and low risk groups, based on the median risk score. Results are illustrated for a discovery cohort (left) and validation cohort (right). (C) Kaplan-Meier analysis of recurrence-free survival for patients based on risk score stratification in a discovery and independent validation cohort as described in (B). High and low risk groups were compared using a standard log-rank test.
